# Adaptation and convergence in circadian-related genes in Iberian freshwater fish

**DOI:** 10.1101/706713

**Authors:** João M Moreno, Tiago F Jesus, Vitor C Sousa, Maria M Coelho

## Abstract

**Background:** The circadian clock is a biological timing system that improves the ability of organisms to deal with environmental fluctuations. At the molecular level it consists of a network of transcription-translation feedback loops, involving genes that activate (*bmal* and *clock* – positive loop) and repress expression (cryptochrome (*cry*) and period (*per*) – negative loop). This is regulated by daily alternations of light but can also be affected by temperature. Fish, as ectothermic, depend on the environmental temperature and thus are good models to study its integration within the circadian system. Here, we studied the molecular evolution of circadian genes in four *Squalius* freshwater fish species, distributed across Western Iberian rivers affected by two climatic types with different environmental conditions (e.g. light and temperature). *S. carolitertii* and *S. pyrenaicus* inhabit the colder northern region under Atlantic climate type, while *S. torgalensis, S. aradensis* and some populations of *S. pyrenaicus* inhabit the warmer southern region affected by summer droughts, under Mediterranean climate type.

**Results:** We identified 16 circadian-core genes in the *Squalius* species using a comparative transcriptomics approach. We detected evidence of positive selection in nine of these genes using methods based on dN/dS. Positive selection was mainly found in *cry* and *per* genes of the negative loop of the cycle, with 11 putatively adaptive substitutions mostly located on protein domains. Evidence for positive selection is predominant in southern populations affected by the Mediterranean climate type. By predicting protein features we found that changes at sites under positive selection can impact protein thermostability by changing their aliphatic index and isoelectric point. Additionally, in nine genes, the phylogenetic clustering of species that belong to different clades but inhabit southern basins with similar environmental conditions indicated evolutionary convergence.

**Conclusions:** Our results support that temperature may be a strong selective pressure driving the evolution of genes involved in the circadian system. By integrating sequence-based functional protein prediction with dN/dS-based methods to detect selection we also uncovered adaptive convergence in the southern populations, probably related to their similar thermal conditions.

## BACKGROUNG

Organisms are exposed to daily environmental fluctuations in their natural habitats. To overcome them, organisms developed biological timing systems to optimize their physiological and biochemical processes in space and time [1]. These systems work as internal clocks and require a proper synchronization with environmental signals. Thus, understanding the molecular evolution of the genes involved in the circadian system provide important clues to elucidate how species adapt to their environments.

The circadian system is a universal biological timing system found virtually in all organisms [2]. Circadian system is synchronised by light–dark cycle of a day’s period and present oscillations with a period of ∼24h called circadian rhythms. Oscillations are generated and regulated at molecular level, but the outcomes have been shown to influence several aspects of physiology, behaviour and ecology of organisms [2, 3]. In fact, circadian rhythms have been shown to improve the inherent ability of several organisms to survive under changing environments, by aiding them to efficiently anticipate periodic events, specifically light changes and climate seasons [2, 3].

The molecular circadian system consists of a network of signalling transduction pathways regulated mainly by interconnected transcription-translation feedback loops (Fig. S1) [4, 5]. The regulatory loops are sustained by the so-called core circadian genes and proteins, requiring about 24h to complete a cycle [1, 5]. In vertebrates, several genes have been reported to be responsible for the maintenance and regulation of the circadian system [1]. The core circadian-genes belong to four main gene families: Cryptochromes (CRY), Period (PER), CLOCK, and BMAL [5]. These gene families encompass several characterized genes (*cry, per, bmal* and *clock*) in vertebrates. In fish, several families possess a larger number of circadian paralogs as compared to other vertebrates [6]. For instance, in zebrafish (*Danio rerio*), several genes have been identified: six *cry* (*cry1aa, cry1ab, cry1ba, cry1bb, cry2, cry3*), four *per* (*per1a, per1b, per2, per3*), three *bmal* (*bmal1a, bmal1b, bmal2*), and three *clock* (*clocka, clockb, clock2/npas2*) [7–10]. Cryptochrome genes encode for a class of flavoproteins that are sensitive to blue light [10], whereas *period* genes encode for proteins that also display a strong but differential light responsiveness [11, 12]. Both *cry* and *per* were found to be key agents in the entrainment of the circadian system, as they constitute the negative elements of the system (i.e. repressors of transcription) [13]. BMAL (Brain and muscle ARNT like) and CLOCK (Circadian locomotor output cycle kaput) families encode for canonical circadian proteins, a highly conserved bHLH (basic-Helix-Loop-Helix)-PAS (Period-Aryl hydrocarbon receptor nuclear translocator-Single minded) transcriptional factors and are the positive elements of the circadian system (i.e. activators of transcription) [1, 5].

Studies in fish allowed to elucidate the different levels of organization of the circadian system. This is because fish are a very diverse group of animals adapted to nearly all aquatic environments and possess a larger number of circadian paralogs as compared to the other vertebrates [6]. Advances in genome sequencing allowed to identify circadian-related genes in several model organisms, including zebrafish.

Homology-based methods allowed the identification of circadian genes in other non-model fish species [7–9, 14]. Some studies cover the evolutionary relationships of the core-clock gene families and the mechanisms driving their molecular evolution [7–10, 15], but several key questions remain open, namely the higher number of paralogs in fish when compared to other vertebrates, and its importance for adaptation of species to different environments [6].

By sequencing the transcriptome of zebrafish exposed to light, two recent studies identified several genes whose expression depends on light, revealing a multi-level regulation of circadian rhythms by light-cycles [16, 17]. Photoreception is particularly interesting in fish as, contrary to most vertebrates that only perceive light through the eyes, fish also possess a photosensitive pineal gland, dermal melanophores, and brain photoreceptors [1]. In addition, fish possess independent peripheral photoreceptors and self-sustaining circadian oscillators in every tissue [1, 18].

Circadian rhythms can also be entrained by temperature [19–22]. In mammals it was demonstrated that peripheral cells *in vitro* could sense the change of room temperature as a cue for entrainment of circadian system [19]. In zebrafish, temperature has also an important role in circadian clock [20, 22], and it was proposed that temperature could entrain the phase of the system by driving expression levels of *per3*, and other circadian genes (namely *cry2* and *cry1ba*) via an alternative hypothetical enhancer [20]. In this model, *per1b* (formerly known as *per4*) promoter integrates temperature and light regulatory inputs [20]. In agreement with the hypothesis that temperature affects the circadian system, in a study comparing a transcriptome profiling of two freshwater fish species (*Squalius carolitertii* and *S. torgalensis*) exposed to different temperatures, Jesus et al. [23, 24] found two differentially expressed genes (*cry1aa* and *per1a*) between a control and a thermal stress condition.

In the Western Iberian Peninsula, there are four known species of the genus *Squalius* Bonaparte, 1837 in Portuguese rivers (*S. carolitertii, S. pyrenaicus, S. torgalensis* and *S. aradensis*, Fig. 1). *S. carolitertii* (Doadrio 1988) inhabits the northern river basins, *S. pyrenaicus* (Günther 1868) occurs in the Central and Southern basins (e.g. Tagus, Guadiana and Almargem), while sister species *S. torgalensis* and *S. aradensis* (Coelho et al. 1998) are confined to small Southwestern basins (e.g. Mira and Arade, respectively). Based on phylogenies of nuclear and mitochondrial markers, the species tree of these species is well known, comprising two main groups: (i) *S. carolitertii* and *S. pyrenaicus*; and (ii) *S. torgalensis* and *S. aradensis* [25, 26]. In Western Iberian Peninsula there is a transition between two contrasting climate types (Fig. 1): the Atlantic in the Northern region that is characterized by mild temperatures (inhabited by *S. carolitertii* and *S. pyrenaicus*), and the Mediterranean in the Southern region (inhabited by *S. pyrenaicus, S. torgalensis* and *S. aradensis*) typified by higher temperatures and droughts during summer periods [27–29]. Thus, species inhabiting the southern basins affected by the Mediterranean climate face harsher conditions. Besides differences in spring average water temperature of approximately 5 °C along the distribution of these species, there are differences in the average spring photoperiod (approximately 15 minutes) between the northern and southern basins. The environmental differences associated with the distribution of *Squalius* species in Portugal make them a good model to study adaptation to different environmental conditions.

**Fig. 1.**
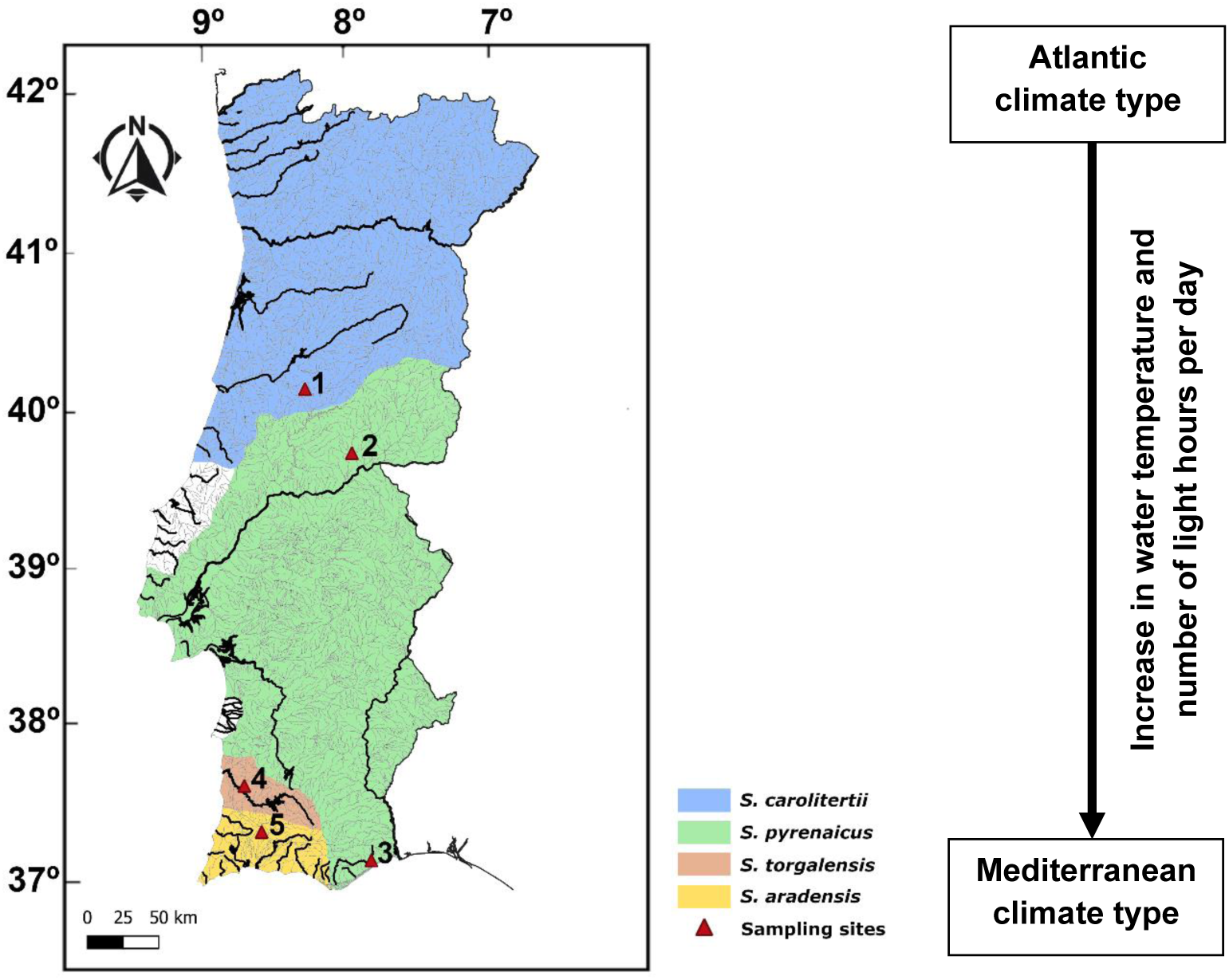
Spatial distribution of the four Portuguese *Squalius* species. Sampling sites are marked with red triangles: Mondego basin (**1**, Sótão river); Tagus basin (**2**, Ocreza river); Almargem basin (**3**, Almargem stream); Mira basin (**4**, Torgal stream); Arade basin (**5**, Odelouca stream). Axes on map represent latitude and longitude, respectively.

Here, we performed an integrative study on the molecular evolution of circadian system genes in *Squalius* species. We aimed to identify the genes involved in the core circadian system in these species and assess their evolutionary history. We combined several approaches, conducting phylogenetic analysis of the identified genes within each gene family in the *Squalius* species and using predicted protein sequences. Finally, we aimed to detect signatures of positive selection and correlate those with predicted functional features of the proteins potentially relevant for the response to environmental differences in light and temperature. These results contribute to a better understanding of the mechanisms of adaptation and response of freshwater fish species to climate change.

## RESULTS

### (i) Evolutionary history of circadian-related gene families

To identify the genes involved in the central circadian system in *Squalius* species, we compared published transcriptome data of *S. torgalensis* and *S. carolitertii* [30] with light-induced zebrafish transcriptomes [16, 17], as these allow to identify more easily genes from circadian systems with relation with light stimuli. From this comparison we identified sixteen genes in the *Squalius* genus belonging to four main gene families (Cryptochromes, Period, CLOCK and BMAL), involved in the main core of circadian system (Table S1) and already described for other fish species [7–10]. Identified genes were re-sequenced for *S. torgalensis* and *S. carolitertii* and sequenced *de novo* for *S. pyrenaicus* and *S. aradensis*, and exon identity was confirmed based on sequence alignment with zebrafish sequences (Table S2).

To test if the proteins encoded by the identified genes potentially retain the function involved in the circadian system, we used the predicted protein sequences to conduct an analysis of protein-protein interactions (PPI). This revealed that all proteins interact with other core circadian proteins (Table S3). We also found proteins with interactions indirectly related with the core circadian system through circadian-dependent functions (Table S3).

To assess evolutionary history of gene families we conducted phylogenetic analysis independently for each family based on predicted protein sequences (Fig. S2). To account for paralogous, we used *Drosophila melanogaster* protein as outgroup and included zebrafish (*Danio rerio*) proteins in the phylogeny (see Table S2 for accession numbers). As detailed for each family below, all reconstructed phylogenies recovered the paralogous evolutionary relationships previously described for other fish species (Fig. S2) [7–10]. We found that *Squalius* sequences form monophyletic groups clustering with orthologs from zebrafish for all gene families. Therefore, our phylogenies support the identification of orthologs in each gene family.

The gene trees based on nucleotide sequences can be grouped in three major topologies: (1) congruent with the species tree, with two main clusters: i) *S. aradensis* and *S. torgalensis*; ii) *S. pyrenaicus* and *S. carolitertii*; (2) incongruent with species tree and with two main clustering groups: i) *S. carolitertii* and *S. pyrenaicus* from Tagus; and ii) *S. torgalensis, S. aradensis* and *S. pyrenaicus* from Almargem; and (3) incongruent with the species tree and with two main clustering groups: i) *S. carolitertii, S. pyrenaicus* from Tagus and S. *torgalensis*; and ii) *S. aradensis* and *S. pyrenaicus* from Almargem (Table 1, Fig. S3). Three additional topologies were recovered for three genes, likely resulting from incomplete lineage sorting (Table 1, Fig. S3).

**Table 1.** Topology of gene trees and summary of tests for positive selection.

| Topology of phylogenetic tree | Genes with topology | Gene-wide test for positive selection | Branch test for positive selection |  | Site-level positive selection |  |
| --- | --- | --- | --- | --- | --- | --- |
|  |  | Positive selection | Positive selection | Branch of the phylogeny under selection | Positive selection (Number of sites) | Details on mutation under selection |
| Species tree<br>((Sc,SpT),SpG),(Sa,St)* | *cry1ba* | ✗ | ✗ | – | ✗ (-) | – |
|  | *per1a* | ✗ | ✗ | – | ✓ (1) | Table 2 |
|  | *per3* | ✓ | ✓ | *S. pyrenaicus*<br>(Almargem) | ✓ (1) | Table 2 |
|  | *bmal2* | ✓ | ✗ | – | ✓ (1) | Table 2 |
| Convergence<br>(Sc,SpT),(St,(Sa,SpA))* | *cry1aa* | ✗ | ✗ | – | ✓ (1) | Table 2 |
|  | *cry1bb* | ✗ | ✗ | – | ✗ (-) | – |
|  | *clockb* | ✓ | ✓ | *S. carolitertii* | ✗ (-) | – |
|  | *clock2* | ✗ | ✗ | – | ✗ (-) | – |
| Convergence<br>((Sc,SpT),St),(Sa,SpA)* | *cry3* | ✗ | ✗ | – | ✓ (1) | Table 2 |
|  | *per1b* | ✓ | ✓ | *S. torgalensis* | ✓ (1) | Table 2 |
|  | *per2* | ✓ | ✓ | *S. torgalensis* + *S. aradensis* | ✓ (4) | Table 2 |
|  | *clocka* | ✗ | ✗ | – | ✗ (-) | – |
|  | *bmal1a* | ✗ | ✗ | – | ✗ (-) | – |
| Other topologies | *cry1ab* | ✗ | ✗ | – | ✗ (-) | – |
|  | *cry2* | ✗ | ✗ | – | ✓ (1) | Table 2 |
|  | *bmal1b* | ✗ | ✗ | – | ✗ (-) | – |

\*Sc, *Squalius carolitertii*; SpT, *Squalius pyrenaicus* (Tagus population); SpA, *Squalius pyrenaicus* (Almargem population); St, *Squalius torgalensis*; Sa, *Squalius aradensis*.

### (ii) Signatures of selection in circadian-related genes

Given the confidence on gene identity, we tested these genes for the presence of signatures of natural selection with several statistical tests based on dN/dS ratio. Out of sixteen genes, we detected positive selection in five using the gene-wide level test implemented in BUSTED (Table 1; Table S4), and in eight using the site-level analysis implemented in MEME to detect individual sites under positive selection (Table 2). Among the genes with signatures of positive selection for at least one of the mentioned methods, we detected significant positive selection in at least one external branch of the phylogeny in five of them, using a branch-based test implemented in aBSREL (Table 1; Table S5).

**Table 2.** Sites under positive selection and protein properties affected.

| Protein | #sites under positive selection | Sites under positive selection | p-value | Type of non-synonymous mutation | Species* | Most probable impacted physicochemical parameter |
| --- | --- | --- | --- | --- | --- | --- |
| **CRY1AA** | 1 | K275Q | 0.03 | Non-conservative | *S. torgalensis* | Isoelectric point (pI) |
| **CRY1AB** | 0 | - | - | - | - | - |
| **CRY1BA** | 0 | - | - | - | - | - |
| **CRY1BB** | 0 | - | - | - | - | - |
| **CRY2** | 1 | I274V | 0.00 | Conservative | *S. aradensis* / *S. torgalensis* | None |
| **CRY3** | 1 | Q96H | 0.02 | Non-conservative | *S. aradensis* / *S. torgalensis* | pI |
| **PER1A** | 1 | T254A | 0.08 | Non-conservative | *S. carolitertii* | Aliphatic index |
| **PER1B** | 1 | T1256V | 0.08 | Non-conservative | *S. torgalensis* | Aliphatic index |
| **PER2** | 4 | T402D | 0.01 | Non-conservative | *S. aradensis* / *S. pyrenaicus* (Almargem) | pI |
|  |  | T514L | 0.02 | Non-conservative | *S. aradensis* | Aliphatic index |
|  |  | R697A | 0.03 | Non-conservative | *S. torgalensis* | pI; Aliphatic index |
|  |  | G1208T | 0.09 | Non-conservative | *S. torgalensis* | None |
| **PER3** | 1 | K429Y | 0.02 | Non-conservative | *S. carolitertii* / *S. pyrenaicus* (Tagus) | pI |
| **CLOCKA** | 0 | - | - | - | - | - |
| **CLOCKB** | 0 | - | - | - | - | - |
| **CLOCK2** | 0 | - | - | - | - | - |
| **BMAL1A** | 0 | - | - | - | - | - |
| **BMAL1B** | 0 | - | - | - | - | - |
| **BMAL2** | 1 | R20P | 0.09 | Non-conservative | *S. torgalensis* | pI |

\*The species/populations with the mutated form was determined according to the comparison with the amino acid present in the reference (*D. rerio*) sequence.

Additionally, we tested for purifying selection with the FEL statistical test to detect sites under negative selection. We found a considerable amount of conserved positions under negative selection, ranging from 2 sites in *cry1ab* to 59 sites in *per2* (Table S6). Moreover, a high proportion of these sites are in codons for aliphatic amino acids (alanine, isoleucine, leucine, proline, and valine) (Table S6).

### (iii) Prediction of functional and structural features of circadian-related proteins

To investigate possible functional modifications on the proteins related with adaptive changes, we predicted the aliphatic index (AI) and isoelectric point (pI) for these proteins based on predicted amino acid sequences. These parameters were chosen due to their relationship with protein thermostability, a factor with high biological relevance for the studied species, given that they inhabit regions with distinct water temperatures. While AI is defined as the relative volume occupied by aliphatic side chains (Ala, Val, Ile, Leu) and it is positively correlated with increased thermostability of globular proteins [31, 32]; pI reflects the pH at which a protein has neutral net charge, which is indicative of the overall composition of charged amino acids, and hence it is important for protein subcellular localization and potential electrostatic interactions responsible from protein stabilization [32–34].

Regarding the predicted aliphatic index (AI), we found differences among *Squalius* species for all proteins except for BMAL1B. For CRY1AA, CRY1AB and PER1A we found that northern populations of *S. carolitertii* and *S. pyrenaicus* from Tagus presented higher values of AI, whereas for CRY3, PER2, PER3, CLOCKB and BMAL1A the southern populations showed higher AI (Table S7). We found differences in the predicted pI across *Squalius* species for most proteins, except for PER1B, CLOCKA, BMAL1A, BMAL1B (Table S7). The pI values tended to be lower in for most proteins of southern populations of *S. torgalensis, S. aradensis* and *S. pyrenaicus* from Almargem (Table S7), even though this was not the case for CRY1BA, CRY1BB, CRY3, PER3, CLOCKB, CLOCK2 and BMAL2.

We also conducted a sequence-based analysis to predict domain and/or motif location using HMM-based methods (Fig. 2). For CRY proteins, we found two common domains to all CRY proteins: the DNA photolyase domain and the FAD-binding domain. Both domains are responsible for binding chromophores and may be of extreme importance for the activity of these proteins [10]. We found two domains common to all PER proteins: Period-Arnt-Sim (PAS_3/PAS_11) domain and the Period protein 2/3C-terminal region. Additionally, PER1A, PER2 and PER3 present an additional PAS domain relatively well conserved. PAS domains are of extreme importance for PER function as they serve for protein dimerization, but also display activity of photoreception [35, 36]. For CLOCK and BMAL proteins, we found three domains conserved in all two proteins: two Period-Arnt-Sim (PAS fold and PAS_11) domain and the basic helix-loop-helix (bHLH). The bHLH is a protein structural motif that characterizes one of the largest families of dimerizing transcription factors and consists in a DNA-binding region.

**Fig. 2.**
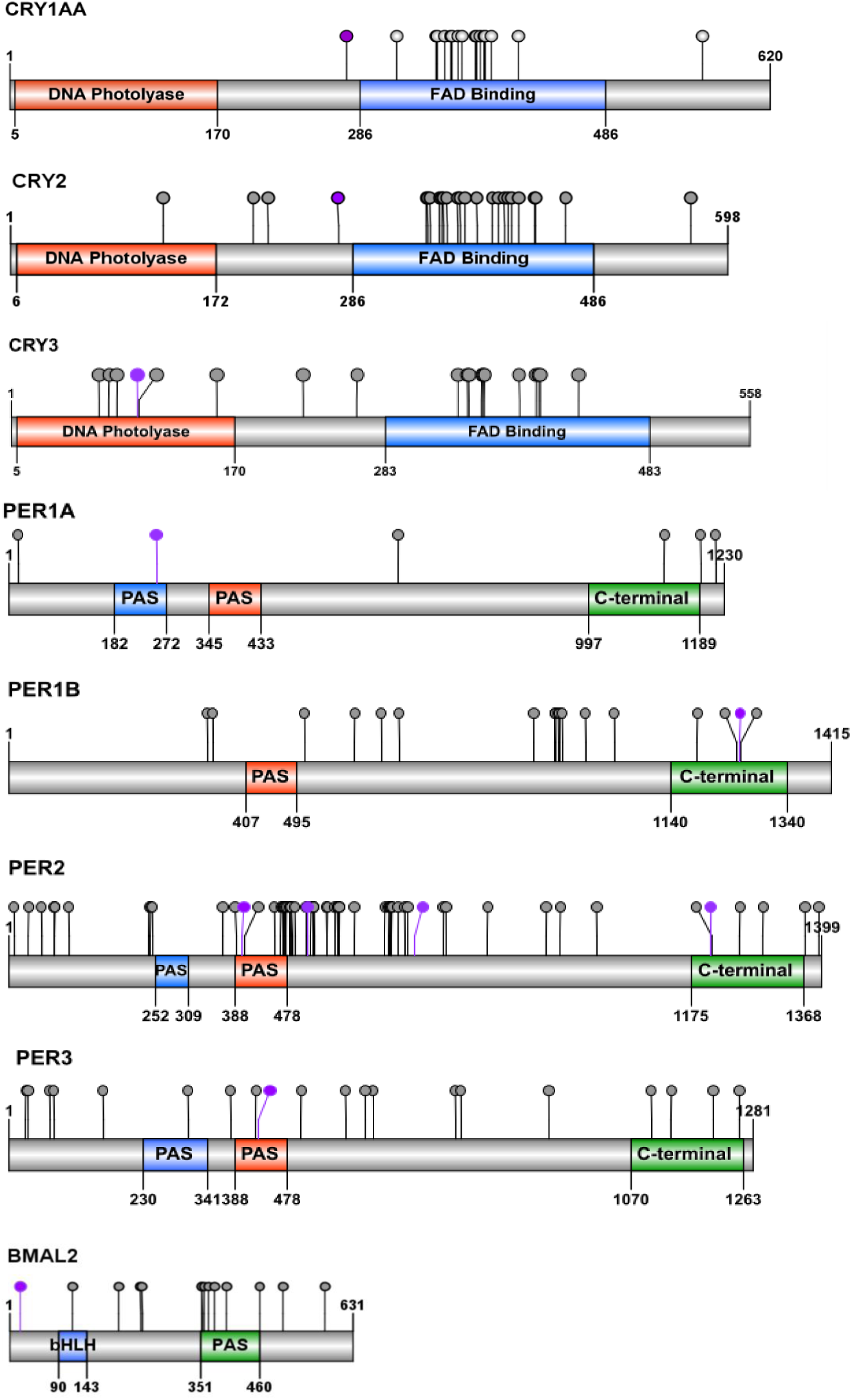
Domain organization of proteins with positively selected sites located inside functional domains; purple dots represent the positively selected positions and grey dots represent negatively selected positions. See Table 2 for further information on sites under positive selection and Table S6 for sites under negative selection.

## DISCUSSION

The circadian system generates and maintains endogenous rhythms synchronised with the daily fluctuation of light and is observed in a wide range of life forms [2, 3]. We identified and characterized the molecular evolution of genes from the core circadian system in western Iberian chubs, freshwater fish species of genus *Squalius* that inhabit different river basins under two climatic types (Atlantic and Mediterranean), with differences in light and temperature.

### Identification of paralogs and orthologs of all circadian gene families in *Squalius*

We were able to identify sixteen genes (Table S1) belonging to the four known core-circadian gene families (CRY, PER, CLOCK and BMAL), which are orthologous to genes described in *D. rerio* and other fish species [6–10].

Phylogenetic relationships within each gene family revealed that the history of paralogs genes is conserved in *Squalius* species, and supports the correct identification of circadian orthologs and paralogs [6]. Moreover, in agreement with results from *Danio rerio* and other fish species [20, 21, 37, 38] (summary on Table S1), we detected possible diversification in functions of the circadian genes that may have arisen to optimize important biological processes, synchronised with the circadian oscillation. We found that in *Squalius* the set of predicted protein-protein interactions in CRY, PER and BMAL (Table S3) involve circadian related proteins, but also proteins with other biological functions, namely BHLHe41 (basic helix-loop-helix family, member e41) and HSF2 (heat-shock factor 2), involved in temperature response [22, 39–41], or with NFIL3-5 (nuclear factor, interleukin 3-regulates, member 5), a protein involved in immune response [42].

### Evidence of positive selection is related with protein predicted function

We found evidence of positive selection mostly on *cry* (*cry1aa, cry2* and *cry3*) and *per* (*per1a, per1b, per2* and *per3*) genes (Tables 1 and 2). These genes encode the negative elements of the circadian system acting as repressors of transcription [43] (Fig. S1). These genes are sensitive to environmental stimuli (e.g. light or temperature) and refine the regulation of the circadian system (e.g. [13, 21, 22]). Moreover, within these genes, we found 11 potential adaptive changes mostly located within the functional domains of the protein, namely in CRY3, PER1A, PER1B, PER2 and PER3 (Table 2, Fig. 2). In CRY3 the most predominant change is located inside the DNA Photolyase domain, one of the light-sensitive domains of cryptochromes. In PER proteins, adaptive substitutions (Table 2) are mostly located inside PAS domains (Fig. 2) that serve for protein dimerization with CRY proteins [35, 36]. Changes in PER2 (T402D) and in PER3 (K429Y) are non-conservative changes that alter the charge of the amino acid at that position, which can consequently alter the strength of protein-protein interaction. Moreover, these changes can impact protein structure as certain amino acids present a propensity for specific structural arrangements [44]. The fact that we find positive selection in the negative elements is in line with previous studies showing that mutations in PER2 protein were important for the adaptation of blind cavefish to its environment [45]. Taken together, these results suggest that circadian system genes were involved in Iberian *Squalius* species adaptation, which occurred mostly by changes in the negative elements of the circadian system.

### More genes under positive selection in populations under Mediterranean climate

Based on dN/dS we found evidence of positive selection in all species using different tests (Tables 1 and 2, Tables S4 and S5). The site level tests indicate that signatures of positive selection are present mostly in southern populations (*S. torgalensis, S. aradensis* and *S. pyrenaicus* from Almargem, Figure 1 and Tables 1 and 2) that are under the influence of the Mediterranean climate type. These results indicate that circadian genes can be involved in adaptation of these species to their specific environments. The circadian system is indirectly related to regulation of many physiological and metabolic aspects that affect the response of organisms to environmental stimuli, and hence these signatures of positive selection can be related to adaptation due to several environmental factors. Despite light stimuli, temperature is important for the proper maintenance of the circadian system in fish [20–22], and temperature is known to impose strong selective pressures in ectothermic species [46, 47]. Thus, present-day and past differences in temperature between river drainage systems inhabited by *Squalius* species can explain our results.

There are several lines of evidence supporting the hypothesis that temperature is a key selective pressure. First, populations with more genes with signatures of positive selection inhabit a region influenced by a Mediterranean climate with higher water temperatures. Second, evidence from protein analysis shows differences in predicted protein thermostability between species in different climatic types. A higher protein thermostability can be achieved either by (i) increasing the aliphatic index AI [31, 32], (ii) increasing the strength of ionic interactions [48], or a combination of these mechanisms. We found that in 11 putatively adaptive changes of the core circadian genes, four of them (in PER1A, PER1B and PER2) have a potential impact on protein aliphatic index (AI), and six of them (in CRY3, PER2, PER3 and BMAL2) on isoelectric point (pI), and therefore can have a direct effect on protein thermostability (Table 2). For these proteins, the analysis of predicted AI and pI showed differences between the *Squalius* species inhabiting the Atlantic and Mediterranean climate types (Tables S7), suggesting that differences in protein thermostability can result from adaptation to different water temperatures. Moreover, the sites inferred to be under negative selection were mostly on codons encoding for aliphatic amino acids, which are associated with protein thermostability.

Last, signatures of positive selection were found mostly on *cry* and *per* genes, circadian genes that have been shown to regulate temperature integration within the circadian system in fish (Table S1) [20, 21, 23, 24]. For instance, we found signatures of positive selection in *cry2, per1b* and *per3*, which are three genes known for integrating temperature within the circadian system in *D. rerio* (Table S1) (see also [20]). A clear example is *per1b*, for which *S. torgalensis* has the PER1B with the highest AI and pI (Table S7) and we found signatures of positive selection in *S. torgalensis* (Table 1) at a site associated with amino changes that increase AI (Table 2), suggesting that selection led to increased thermostability. This protein has been shown to be important in *D. rerio* for the integration of temperature and light cues within the circadian system [20]. We also found signatures of positive selection in *cry1aa* and *per1a*, both found to change their gene expression in *S. torgalensis* and *S. carolitertii* when exposed to increased water temperature under controlled laboratory conditions [23, 24]. In PER1A, we found a mutation under positive selection in *S. carolitertii* (Table 2) at an important region of the protein (PAS domain, Fig. 2) that leads to an increase in the aliphatic index, and therefore increased protein thermostability. This is compatible with Jesus et al. 2017 [24], that observed a downregulation of the expression of this gene at higher temperatures in *S. carolitertii*. We speculate that such increase in protein thermostability could have been selected to properly function at higher temperatures with low expression levels, compensating the costs of over-expression of other proteins.

### Evidence of adaptive convergence from phylogenetic trees and predicted protein function

In four genes (*cry1ba, per1a, per3* and *bmal2*) the best topology for the inferred gene tree is congruent with the species tree with two main clusters: i) *S. aradensis* and *S. torgalensis*; ii) *S. pyrenaicus* and *S. carolitertii* [25, 26]. This could indicate that these genes evolved neutrally during speciation. However, we found signatures of positive selection in three of them (*per1a, per3* and *bmal2*), which might indicate that the evolution of these genes has been at least partially driven by natural selection, rather than exclusively by neutral divergence after speciation events. However, the phylogenies we inferred using both the nucleotide and protein sequences indicate that for 9 out of 16 genes (*cry1aa, cry1bb, cry3, per1b, per2, clockb, clock2* and *bmal1a*) the best topology clusters together, with high support, *S. aradensis* and *S. pyrenaicus* from Almargem (Fig. S2 and S3). According to the species tree these two species belong to two highly divergent lineages [25], and hence such a high proportion of genes with this clustering is unlikely due to neutral incomplete lineage sorting. Instead, this suggests a scenario of evolutionary convergence of populations inhabiting similar environments. In fact, Almargem and Arade are two basins from the south of Portugal influenced by Mediterranean climate, facing very similar environmental conditions (e.g. average water temperature and photoperiod). This pattern of sequence convergence in these genes and proteins may thus be a consequence of convergent adaptation.

The convergence in these genes and proteins matches at least partially the criteria for detecting evolutionary convergence proposed by Dávalos et al. (2012), namely: (1) evidence from sequences of functional parts of genes; (2) clear link between gene function and ecological conditions; and (3) evidence that selection is acting on target genes at different rates from other lineages. As we only obtained data from cDNA, we expect all the sequences to be from exons, and therefore, constituting functional parts of genes, confirming the criterium 1. As previously mentioned, most genes presenting this signal of convergence belong to CRY and PER family, and both *cry* and *per* genes have a demonstrated importance in the response to environmental temperature, therefore confirming the criterium 2. Based on dN/dS tests we could confirm the criterium 3 for *cry3* and *per2*, since at least one of the populations *S. aradensis* or *S. pyrenaicus* (Almargem) show signatures of positive selection. For *cry1aa, per1b* and *clockb*, protein analysis supports a strong similarity between the physicochemical parameters estimated for *S. aradensis* and *S. pyrenaicus* from Almargem, pointing to a functional convergence at protein level, even though for those genes signatures of positive selection were on either *S. torgalensis* or *S. carolitertii*. For *cry1bb, bmal1a* and *clock2* we did not find any evidence for signatures of selection, but for BMAL1A and CLOCK2 proteins, the functional characterisation also revealed similar predicted protein physicochemical patterns in *S. aradensis* and *S. pyrenaicus* from Almargem, pointing to functional convergence. The exception was the *cry1bb* for which we do not have support for convergence at any level except the phylogenetic, and therefore it could be explained by a scenario of incomplete lineage sorting.

Scenarios of convergence have been described for other species at the morphological level [50–52], and at the molecular level [53–58]. Moreover, light was shown to be an important determinant in two studies that detected molecular convergence in fish, such as in: (1) the evolution of albinism linked to the *Oca2* gene in two independent populations of the cavefish *Astyanax mexicanus* [53]; and (2) in functional evolution of Rhodopsin proteins in several fish species [59]. Here, we detected convergent evolution in freshwater fish in genes and proteins related to integration of visual and thermal stimuli within the circadian system, which are part of gene families with duplications. This raises the possibility that gene duplications can be important in convergent evolution, which could be further studied and tested in the future.

## CONCLUSIONS

This study aimed to characterise the evolution of circadian-related gene families in non-model freshwater organisms. These results provide clear insights on how the environment can shape the evolution of the circadian system, and how it can contribute to the process of adaptation to different environments.

Our findings support that together with neutral historical factors, natural selection also drove the molecular evolution of the four studied species that live in different environmental conditions of light and temperature, affected by Atlantic and Mediterranean climate types. Moreover, we find evidence for adaptive convergence between the two southern populations of *S. aradensis* and *S. pyrenaicus* from Almargem, a pattern described for the first time. Our results using an approach combining dN/dS and protein analysis can help understanding the genetic patterns found in other species, namely those that are due to convergence, a process that can be frequent but difficult to detect.

## METHODS

### Sampling

Muscle tissue from five wild adult fish of *S. carolitertii* and *S. torgalensis* species was available from previous work [24] and stored at −80°C in RNALater® (Ambion, Austin, TX, USA). The adult fish were sampled in Portuguese basins of Mondego (40°8’5.22”N; 8°8’35.06”W) and Mira (37°38’1.31”N; 8°37’22.37”W), respectively [24]. Samples of *S. pyrenaicus* were also stored at −80°C from previous projects. This work includes two sampling sites: Almargem (37°09’50.7”N; 7°37’13.2”W) [60] and Tagus (39°43’48.2”N; 7°45’38.1”W) [61].

*Squalius aradensis* individuals were captured from Portuguese basin Arade (37°17’0.53’’N; 8°29’7.31’’W) under the license 421/2017/CAPT issued by Portuguese authority for Conservation of endangered species [ICNF (Instituto da Conservação da Natureza e das Florestas)]. After capturing, the specimens were transported alive to the laboratory in aerated containers to minimize fish discomfort and euthanized immediately upon arrival with an overdose of tricaine mesylate (400 ppm of MS-222; Sigma-Aldrich, St. Louis, MO, USA) with sodium bicarbonate (1:2) following the recommended ethical guidelines (ASAB/ABS, 2012) and European Union regulations. Organs were stored in RNALater® at −80°C until further use. Distribution of the species and sampling sites are illustrated in Fig. 1.

### Identification of predicted circadian system related genes in Iberian freshwater fish based on transcriptome analysis

Previously published transcriptome assemblies of *S. torgalensis* and *S. carolitertii* obtained from RNA-Seq experiments [30], were used to identify the genes related to the circadian system in the study species. To identify potential genes related to the circadian system we performed BLAST searches of the transcriptomes of the two species against two *Danio rerio* light-induced transcriptomes [16, 17]. To account for splicing isoforms and to avoid the misidentification of potential paralogous genes, we used a stringent e-value threshold of 1×10^−7^ for the BLAST searches and kept only sequences with identity higher than 85%. To predict the biological and molecular function of the identified genes, we performed a functional enrichment analysis using the list of top blast *Danio rerio* ENA accession numbers (Table S2) and the method implemented in DAVID functional annotation tool. Through this methodology we were able to find enriched GO terms among the genes retrieved. The most significant enriched GO terms for Biological Process and Molecular Function were filtered, and only the core genes from circadian system (i.e. those described to be involved in the feedback loop) were further kept (Table S1). A threshold for adjusted p-values (Benjamini) of 0.05 was used to remove false positives.

### Gene sequencing and protein sequence prediction

Based on the sequences retrieved from the transcriptomes of *S. torgalensis* and *S. carolitertii* that matched core genes of the circadian system, we designed specific primers for Polymerase chain reactions (PCRs) using PerlPrimer software v.1.1.19 [62] (Table S8) with the purpose of amplifying the coding regions of those same genes for all the studied species. We included *D. rerio* sequences (Table S2) during primer design to identify conserved regions among Cyprinidae. To further distinguish orthologs from paralogs we performed phylogenetic analysis (see below).

Total RNA was extracted from muscle samples of 25 individuals, 5 from each population. RNA was used to facilitate gene sequencing since introns are avoided. Only muscle tissue was used to avoid tissue expression bias. We added 1 mL TRI Reagent (Ambion, Austin, TX, USA) to 50–100 mg of muscle samples and, after homogenization with Tissue Ruptor (Qiagen, Valencia, CA, USA), extracted RNA according to the TRI Reagent manufacturers protocol. TURBO DNase (Ambion, Austin, TX, USA) was employed to degrade any remaining genomic DNA contaminants, followed by phenol/chloroform purification and LiCl precipitation [63]. Sample quality was checked using a NanoDrop™-1000 spectrophotometer (Thermo Fisher Scientific, Waltham, MA, USA) based on the 260nm/280nm and 260nm/230nm absorbance ratios. Samples concentration was determined with Qubit® 2.0 Fluorometer (Thermo Fisher Scientific, Waltham, MA, USA) to ensure enough quantity of homogeneous RNA for cDNA synthesis. Synthesis of cDNA was performed according to manufacturer’s protocol using the RevertAid H Minus First Strand cDNA synthesis kit (Thermo Fisher Scientific, Waltham, MA, USA), and it was stored subsequently at - 20°C until further use. PCRs were performed in 25 μL reactions containing 10–100 ng of cDNA, 2 mM MgCl_2_, 2 mM each dNTP, 10 μM each primer, Taq Polymerase (5 U/μL), and 1× Taq buffer using thermocycler conditions described in Table S9. PCR products were confirmed using a 1% agarose gel electrophoresis, and after purification with ExoSAP-IT® PCR Product Cleanup (Affymetrix, Inc., Santa Clara, CA, USA), they were sequenced by Sanger sequencing.

Sequences were aligned and edited using Sequencher v.4.2 (Gene Codes Corp., Ann Arbor, MI, USA). Non-redundant nucleotide sequences were deposited in European Nucleotide Archive (ENA) database under the accession numbers available in supplementary Table S2. CLC Sequence Viewer v.7.5. (CLC bio, Aarhus, Denmark) was used to predict protein sequences for *in silico* analysis. BLAST searches were conducted with resulting protein sequences against UniProt database [64] to ensure their identity. Protein sequences for *Danio rerio* were retrieved from UniProt database for each protein (Table S2), as well as *Drosophila melanogaster* homolog sequence for each gene family (Table S2) to use as outgroup in phylogenetic analysis (see below). Protein sequences were aligned by gene family using the M-Coffee method, that combines several alignment algorithms (eg. MUSCLE, MAFFT and CLUSTAL) [65] available in the T-Coffee web server [66]. For each individual gene, nucleotide sequences of *Squalius* species were also aligned using M-Coffee.

### Phylogenetic analysis and molecular evolution

The most appropriate model for amino acid substitution for each data set was determined with ProtTest v.3.0 [67], using both the Akaike information criteria and Bayesian information criteria. Phylogenetic trees were reconstructed for each gene family independently using the protein sequences and the Bayesian Inference method implemented in MrBayes v.3.2.6 [68, 69], using *D. melanogaster* protein sequences (Table S2) as outgroup. The Monte Carlo Markov Chain (MCMC) were ran for 500,000 generations using the parameters determined in ProtTest for protein sequences as priors. Trees were sampled every 500 generations during the analysis. The first 50,000 generations were excluded as burn-in after examining the variation in log-likelihood scores over time. Phylogenetic trees were constructed using protein sequences instead of nucleotide sequences to avoid bias from the third codon rapid evolution. Protein sequences used correspond to the direct translation of nucleotide sequences under the standard genetic code. These phylogenetic trees allowed to confirm the correct identification of orthologs and paralogs.

Gene trees were reconstructed for each gene based on nucleotide sequences using a Maximum Likelihood approach on RAxML v.8 [70] only including *Squalius* sequences. The most appropriate model for nucleotide substitution was determined using MEGA X [71].

All trees were edited in FigTree v1.4.2 (A. Rambaut, University of Edinburgh, UK; http://tree.bio.ed.ac.uk/software/figtree/).

### Analysis of signatures of selection based on dN/dS

Signatures of selection were examined based on the dN/dS ratio (also known as the parameter ω) using four models implemented in HyPhy [72] through the Datamonkey adaptive evolution webserver (Kosakovsky Pond & Frost 2005; Weaver et al. 2018; http://www.datamonkey.org/; accessed in August 2018), including (1) BUSTED (Branch-site Unrestricted Statistical Test for Episodic Diversification) [75] that provides a gene-wide test for positive selection, i.e. it estimates one ω per gene; (2) MEME (Mixed Effects Model of Evolution) [76] a mixed-effects maximum likelihood approach to test the hypothesis that individual sites have been subject to positive selection, i.e. estimating variable ω among sites; (3) aBSREL (adaptive Branch-Site Random Effects Likelihood) [77, 78] for genes whose signals of positive selection were detected with BUSTED or MEME, to test for positive selection on particular branches of the gene tree, i.e. estimating different ω for different branches; and (4) FEL (Fixed Effects Likelihood) [79] that uses a maximum likelihood approach to infer nonsynonymous (dN) and synonymous (dS) substitution rates on a per-site basis for a given coding alignment and corresponding phylogeny and tests the hypothesis that individual sites have been subject to negative selection.

### Prediction of functional and structural features of the predicted proteins

Homology methods available on several resources at the ExPASy Server [32] were used to infer several properties of the predicted proteins from cDNA sequences. Specifically, physicochemical parameters of the proteins were predicted using ProtParam [32]. Given the biological significance of factors related to temperature and pH, we predicted the aliphatic index (AI) and the isoelectric point (pI). We tested for differences in physicochemical parameters using several statistical tests implemented in R v.3.2.3 (R Core Team 2015). First, we checked for normality using the Shapiro-Wilk Test. Due to lack of normality, we used a Kruskal-Wallis Rank Sum Test to identify overall statistical differences in parameters across the populations. When evidence for differences were found, we conducted pairwise Wilcoxon Rank Sum Tests to compare the different groups.

To assess further functional features, a sequence-based prediction of family assignment and sequence domains based on collections of Hidden-Markov Models to support the predictions were accomplished for each protein separately using HMMER web server [80–82] against Pfam database [83]. Representative images of structural organization of domains and locations of sites under selection were created and edited with IBS, Illustrator for Biological Sequences [84].

## DECLARATIONS

### Data accessibility

All sequences obtained during the execution of this work are available on the European Nucleotide Archive (ENA) under the accession numbers reported in Table S1.

### Authors’ contributions

Conceptualization, design and writing by JMM, VCS and MMC. Investigation and data analysis by JMM and TFJ, under supervision of MMC and VCS. Final version approved by all authors.

## Acknowledgements

The authors thank Carla Sousa-Santos for collaborating with fieldwork in collection of specimens of *S. aradensis.* We also acknowledge ICNF (Instituto da Conservação da Natureza e Florestas) for issuing the fishing licenses.

## REFERENCES

1. Foulkes NS, Whitmore D, Vallone D, Bertolucci C. Studying the Evolution of the Vertebrate Circadian Clock. In: Genetics, Genomics and Fish Phenomics. 2016. p. 1–30.

2. Paranjpe DA, Sharma VK. Evolution of temporal order in living organisms. Journal of Circadian Rhythms. 2005;3:7.

3. Vaze KM, Sharma VK. On the adaptive significance of circadian clocks for their owners. Chronobiology international. 2013;30:413–33.

4. Dunlap JC. Molecular Bases for Circadian Clocks. Cell. 1999;96:271–290.

5. Pando MP, Sassone-Corsi P. Unraveling the mechanisms of the vertebrate circadian clock: zebrafish may light the way. BioEssays. 2002;24:419–26.

6. Toloza-Villalobos J, Arroyo JI, Opazo JC. The Circadian Clock of Teleost Fish: A Comparative Analysis Reveals Distinct Fates for Duplicated Genes. Journal of Molecular Evolution. 2015;80:57–64.

7. Wang H. Comparative analysis of period genes in teleost fish genomes. Journal of Molecular Evolution. 2008;67:29–40.

8. Wang H. Comparative analysis of teleost fish genomes reveals preservation of different ancient clock duplicates in different fishes. Marine Genomics. 2008;1:69–78.

9. Wang H. Comparative genomic analysis of teleost fish bmal genes. Genetica. 2009;136:149–161.

10. Liu C, Hu J, Qu C, Wang L, Huang G, Niu P, et al. Molecular evolution and functional divergence of zebrafish (Danio rerio) cryptochrome genes. Scientific Reports. 2015;5:8113.

11. Vatine G, Vallone D, Appelbaum L, Mracek P, Ben-Moshe Z, Lahiri K, et al. Light Directs Zebrafish period2 Expression via Conserved D and E Boxes. PLoS Biology. 2009;7:e1000223.

12. Vallone D, Gondi SB, Whitmore D, Foulkes NS. E-box function in a period gene repressed by light. Proc Natl Acad Sci U S A. 2004;101:4106–11.

13. Tamai TK, Young LC, Whitmore D. Light signaling to the zebrafish circadian clock by Cryptochrome 1a. Proceedings of the National Academy of Sciences of the United States of America. 2007;104:14712–14717.

14. Mei Q, Sadovy Y, Dvornyk V. Molecular evolution of cryptochromes in fishes. Gene. 2015;574:112–20.

15. Sun Y, Liu C, Huang M, Huang J, Liu C, Zhang J, et al. The Molecular Evolution of Circadian Clock Genes in Spotted Gar (Lepisosteus oculatus). Genes. 2019;10:622.

16. Weger BD, Sahinbas M, Otto GW, Mracek P, Armant O, Dolle D, et al. The Light Responsive Transcriptome of the Zebrafish: Function and Regulation. PLoS ONE. 2011;6:e17080.

17. Ben-Moshe Z, Alon S, Mracek P, Faigenbloom L, Tovin A, Vatine GD, et al. The light-induced transcriptome of the zebrafish pineal gland reveals complex regulation of the circadian clockwork by light. Nucleic Acids Research. 2014;42:3750–3767.

18. Whitmore D, Foulkes NS, Strähle U, Sassone-Corsi P. Zebrafish Clock rhythmic expression reveals independent peripheral circadian oscillators. Nature neuroscience. 1998;1:701–707.

19. Tsuchiya Y, Akashi M, Nishida E. Temperature compensation and temperature resetting of circadian rhythms in mammalian cultured fibroblasts. Genes to Cells. 2003;8:713–720.

20. Lahiri K, Vallone D, Gondi SB, Santoriello C, Dickmeis T, Foulkes NS. Temperature Regulates Transcription in the Zebrafish Circadian Clock. PLoS Biology. 2005;3:e351.

21. Chappuis S, Ripperger JA, Schnell A, Rando G, Jud C, Wahli W, et al. Role of the circadian clock gene Per2 in adaptation to cold temperature. Molecular Metabolism. 2013;2:184–93.

22. Jerônimo R, Moraes MN, de Assis LVM, Ramos BC, Rocha T, Castrucci AM de L. Thermal stress in Danio rerio : a link between temperature, light, thermo-TRP channels, and clock genes. Journal of Thermal Biology. 2017;68:128–38.

23. Jesus TF, Grosso AR, Almeida-Val VMF, Coelho MM. Transcriptome profiling of two Iberian freshwater fish exposed to thermal stress. J Therm Biol. 2016;55:54–61.

24. Jesus TF, Moreno JM, Repolho T, Athanasiadis A, Rosa R, Almeida-Val VMF, et al. Protein analysis and gene expression indicate differential vulnerability of Iberian fish species under a climate change scenario. PLOS ONE. 2017;12:e0181325.

25. Sousa-Santos C, Jesus TF, Fernandes C, Robalo JI, Coelho MM. Fish diversification at the pace of geomorphological changes: evolutionary history of western Iberian Leuciscinae (Teleostei: Leuciscidae) inferred from multilocus sequence data. Molecular Phylogenetics and Evolution. 2019;133:263–85.

26. Waap S, Amaral AR, Gomes B, Coelho MM. Multi-locus species tree of the chub genus Squalius (Leuciscinae: Cyprinidae) from western Iberia: new insights into its evolutionary history. Genetica. 2011;139:1009–18.

27. Mesquita N, Coelho MM. The ichthyofauna of the small Mediterranean-type drainages of Portugal: its importance for conservation. In: Conservation of Freshwater Fishes: Options for the Future. 2002. p. 65–71.

28. Mesquita N, Hänfling B, Carvalho GR, Coelho MM. Phylogeography of the cyprinid Squalius aradensis and implications for conservation of the endemic freshwater fauna of southern Portugal. Molecular Ecology. 2005;14:1939–54.

29. Henriques R, Sousa V, Coelho MM. Migration patterns counteract seasonal isolation of Squalius torgalensis, a critically endangered freshwater fish inhabiting a typical Circum-Mediterranean small drainage. Conserv Genet. 2010;11:1859–70.

30. Jesus TF, Grosso AR, Almeida-Val VMF, Coelho MM. Data from: “Characterization of two Iberian freshwater fish transcriptomes, Squalius carolitertii and Squalius torgalensis, living in distinct environmental conditions” in Genomic Resources Notes Accepted 1 April 2015 to 31 May 2015. Molecular Ecology Resources. 2015;16:377.

31. Ikai A. Thermostability and aliphatic index of globular proteins. J Biochem. 1980;88:1895–8.

32. Gasteiger E, Hoogland C, Gattiker A, Duvaud S, Wilkins MR, Appel RD, et al. Protein Identification and Analysis Tools on the ExPASy Server. The Proteomics Protocols Handbook. 2005;:571–607.

33. Kiraga J, Mackiewicz P, Mackiewicz D, Kowalczuk M, Biecek P, Polak N, et al. The relationships between the isoelectric point and: length of proteins, taxonomy and ecology of organisms. BMC Genomics. 2007;8:163.

34. Khaldi N, Shields DC. Shift in the isoelectric-point of milk proteins as a consequence of adaptive divergence between the milks of mammalian species. Biology Direct. 2011;6:40.

35. Ponting CP, Aravind L. PAS: a multifunctional domain family comes to light. Current biology : CB. 1997;7:R674–7.

36. Möglich A, Ayers RA, Moffat K. Structure and Signaling Mechanism of Per-ARNT-Sim Domains. Structure. 2009;17:1282–94.

37. Hirayama J, Cho S, Sassone-Corsi P. Circadian control by the reduction/oxidation pathway: Catalase represses light-dependent clock gene expression in the zebrafish. PNAS. 2007;104:15747–52.

38. Bian S-S, Zheng X-L, Sun H-Q, Chen J-H, Lu Y-L, Liu Y-Q, et al. Clock1a affects mesoderm development and primitive hematopoiesis by regulating Nodal-Smad3 signaling in the zebrafish embryo. J Biol Chem. 2017;292:14165–75.

39. Li Y, Li G, Wang H, Du J, Yan J. Analysis of a Gene Regulatory Cascade Mediating Circadian Rhythm in Zebrafish. PLOS Computational Biology. 2013;9:e1002940.

40. Hung I-C, Hsiao Y-C, Sun HS, Chen T-M, Lee S-J. MicroRNAs regulate gene plasticity during cold shock in zebrafish larvae. BMC Genomics. 2016;17.

41. Tamaru T, Hattori M, Honda K, Benjamin I, Ozawa T, Takamatsu K. Synchronization of Circadian Per2 Rhythms and HSF1-BMAL1:CLOCK Interaction in Mouse Fibroblasts after Short-Term Heat Shock Pulse. PLOS ONE. 2011;6:e24521.

42. Zhang W, Zhang J, Kornuc M, Kwan K, Frank R, Nimer SD. Molecular cloning and characterization of NF-IL3A, a transcriptional activator of the human interleukin-3 promoter. Mol Cell Biol. 1995;15:6055–63.

43. Vatine G, Vallone D, Gothilf Y, Foulkes NS. It’s time to swim! Zebrafish and the circadian clock. FEBS Letters. 2011;585:1485–1494.

44. Chou PY, Fasman GD. Prediction of protein conformation. Biochemistry. 1974;13:222–45.

45. Ceinos RM, Frigato E, Pagano C, Fröhlich N, Negrini P, Cavallari N, et al. Mutations in blind cavefish target the light-regulated circadian clock gene, period 2. Scientific Reports. 2018;8. doi: 10.1038/s41598-018-27080-2.

46. Gunderson AR, Stillman JH. Plasticity in thermal tolerance has limited potential to buffer ectotherms from global warming. Proc Biol Sci. 2015;282:20150401.

47. Paaijmans KP, Heinig RL, Seliga RA, Blanford JI, Blanford S, Murdock CC, et al. Temperature variation makes ectotherms more sensitive to climate change. Global Change Biology. 2013;19:2373–80.

48. Kumar S, Tsai C-J, Nussinov R. Factors enhancing protein thermostability. Protein Engineering, Design and Selection. 2000;13:179–91.

49. Dávalos LM, Cirranello AL, Geisler JH, Simmons NB. Understanding phylogenetic incongruence: lessons from phyllostomid bats. Biological Reviews. 2012;87:991–1024.

50. Muschick M, Indermaur A, Salzburger W. Convergent Evolution within an Adaptive Radiation of Cichlid Fishes. Current Biology. 2012;22:2362–8.

51. Alter SE, Brown B, Stiassny MLJ. Molecular phylogenetics reveals convergent evolution in lower Congo River spiny eels. BMC Evolutionary Biology. 2015;15:224.

52. Passow CN, Arias-Rodriguez L, Tobler M. Convergent evolution of reduced energy demands in extremophile fish. PLoS One. 2017;12.

53. Protas ME, Hersey C, Kochanek D, Zhou Y, Wilkens H, Jeffery WR, et al. Genetic analysis of cavefish reveals molecular convergence in the evolution of albinism. Nature Genetics. 2006;38:107–11.

54. Nath A, Chaube R, Subbiah K. An insight into the molecular basis for convergent evolution in fish antifreeze Proteins. Computers in Biology and Medicine. 2013;43:817–21.

55. Natarajan C, Hoffmann FG, Weber RE, Fago A, Witt CC, Storz JF. Predictable convergence in hemoglobin function has unpredictable molecular underpinnings. Science. 2016;354:336–9.

56. Zhu X, Guan Y, Signore AV, Natarajan C, DuBay SG, Cheng Y, et al. Divergent and parallel routes of biochemical adaptation in high-altitude passerine birds from the Qinghai-Tibet Plateau. Proceedings of the National Academy of Sciences. 2018;115:1865–70.

57. Castiglione GM, Schott RK, Hauser FE, Chang BSW. Convergent selection pressures drive the evolution of rhodopsin kinetics at high altitudes via nonparallel mechanisms. Evolution. 2018;72:170–86.

58. Graham AM, McCracken KG. Convergent evolution on the hypoxia-inducible factor (HIF) pathway genes EGLN1 and EPAS1 in high-altitude ducks. Heredity. 2019. doi: 10.1038/s41437-018-0173-z.

59. Yokoyama S, Tada T, Zhang H, Britt L. Elucidation of phenotypic adaptations: Molecular analyses of dim-light vision proteins in vertebrates. Proceedings of the National Academy of Sciences. 2008;105:13480–5.

60. Machado MP, Matos I, Grosso AR, Schartl M, Coelho MM. Non-canonical expression patterns and evolutionary rates of sex-biased genes in a seasonal fish. Molecular Reproduction and Development. 2016;83:1102–1115.

61. Matos IMN, Coelho MM, Schartl M. Gene copy silencing and DNA methylation in natural and artificially produced allopolyploid fish. The Journal of Experimental Biology. 2016;219:3072–81.

62. Marshall OJ. PerlPrimer: cross-platform, graphical primer design for standard, bisulphite and real-time PCR. Bioinformatics. 2004;20:2471–2.

63. Cathala G, Savouret J-F, Mendez B, West BL, Karin M, Martial JA, et al. A Method for Isolation of Intact, Translationally Active Ribonucleic Acid. DNA. 1983;2:329–35.

64. The UniProt Consortium. UniProt: a hub for protein information. Nucleic Acids Research. 2015;43:D204–12.

65. Wallace IM, O’Sullivan O, Higgins DG, Notredame C. M-Coffee: combining multiple sequence alignment methods with T-Coffee. Nucleic Acids Res. 2006;34:1692–9.

66. Di Tommaso P, Moretti S, Xenarios I, Orobitg M, Montanyola A, Chang J-M, et al. T-Coffee: a web server for the multiple sequence alignment of protein and RNA sequences using structural information and homology extension. Nucleic Acids Research. 2011;39 suppl:W13–W17.

67. Darriba D, Taboada GL, Doallo R, Posada D. ProtTest 3: fast selection of best-fit models of protein evolution. Bioinformatics. 2011;27:1164–5.

68. Huelsenbeck JP, Ronquist F. MRBAYES: Bayesian inference of phylogenetic trees. Bioinformatics. 2001;17:754–755.

69. Ronquist F, Teslenko M, Van Der Mark P, Ayres DL, Darling A, Höhna S, et al. Mrbayes 3.2: Efficient bayesian phylogenetic inference and model choice across a large model space. Systematic Biology. 2012;61:539–542.

70. Stamatakis A. RAxML version 8: a tool for phylogenetic analysis and post-analysis of large phylogenies. Bioinformatics. 2014;30:1312–3.

71. Kumar S, Stecher G, Li M, Knyaz C, Tamura K. MEGA X: Molecular Evolutionary Genetics Analysis across Computing Platforms. Mol Biol Evol. 2018;35:1547–9.

72. Kosakovsky Pond SL, Frost SDW, Muse VS. HyPhy: Hypothesis testing using phylogenies. Bioinformatics. 2005;21:676–679.

73. Kosakovsky Pond SL, Frost SDW. Datamonkey: Rapid detection of selective pressure on individual sites of codon alignments. Bioinformatics. 2005;21:2531–2533.

74. Weaver S, Shank SD, Spielman SJ, Li M, Muse VS, Kosakovsky Pond SL. Datamonkey 2.0: A Modern Web Application for Characterizing Selective and Other Evolutionary Processes. Molecular Biology and Evolution. 2018;35:773–777.

75. Murrell B, Weaver S, Smith MD, Wertheim JO, Murrell S, Aylward A, et al. Gene-wide identification of episodic selection. Molecular Biology and Evolution. 2015;32:1365–1371.

76. Murrell B, Wertheim JO, Moola S, Weighill T, Scheffler K, Kosakovsky Pond SL. Detecting individual sites subject to episodic diversifying selection. PLoS Genetics. 2012;8:e1002764.

77. Kosakovsky Pond SL, Murrell B, Fourment M, Frost SDW, Delport W, Scheffler K. A random effects branch-site model for detecting episodic diversifying selection. Molecular Biology and Evolution. 2011;28:3033–3043.

78. Smith MD, Wertheim JO, Weaver S, Murrell B, Scheffler K, Kosakovsky Pond SL. Less is more: An adaptive branch-site random effects model for efficient detection of episodic diversifying selection. Molecular Biology and Evolution. 2015;32:1342–1353.

79. Kosakovsky Pond SL, Frost SDW. Not so different after all: A comparison of methods for detecting amino acid sites under selection. Molecular Biology and Evolution. 2005;22:1208–1222.

80. Finn RD, Clements J, Eddy SR. HMMER web server: interactive sequence similarity searching. Nucleic Acids Research. 2011;39 suppl:W29–37.

81. Prakash A, Jeffryes M, Bateman A, Finn RD. The HMMER Web Server for Protein Sequence Similarity Search. Current Protocols in Bioinformatics. 2017;60:3.15.1-3.15.23.

82. Potter SC, Luciani A, Eddy SR, Park Y, Lopez R, Finn RD. HMMER web server: 2018 update. Nucleic Acids Res. 2018;46:W200–4.

83. Finn RD, Coggill P, Eberhardt RY, Eddy SR, Mistry J, Mitchell AL, et al. The Pfam protein families database: towards a more sustainable future. Nucleic Acids Research. 2016;44:D279–85.

84. Liu W, Xie Y, Ma J, Luo X, Nie P, Zuo Z, et al. IBS: an illustrator for the presentation and visualization of biological sequences. Bioinformatics. 2015;31:3359–61.

