## Supplementary material for "Adaptation and convergence in circadian-related genes in Iberian freshwater fish": Fig. S

**for**

### SUPPLEMENTARY FIGURES

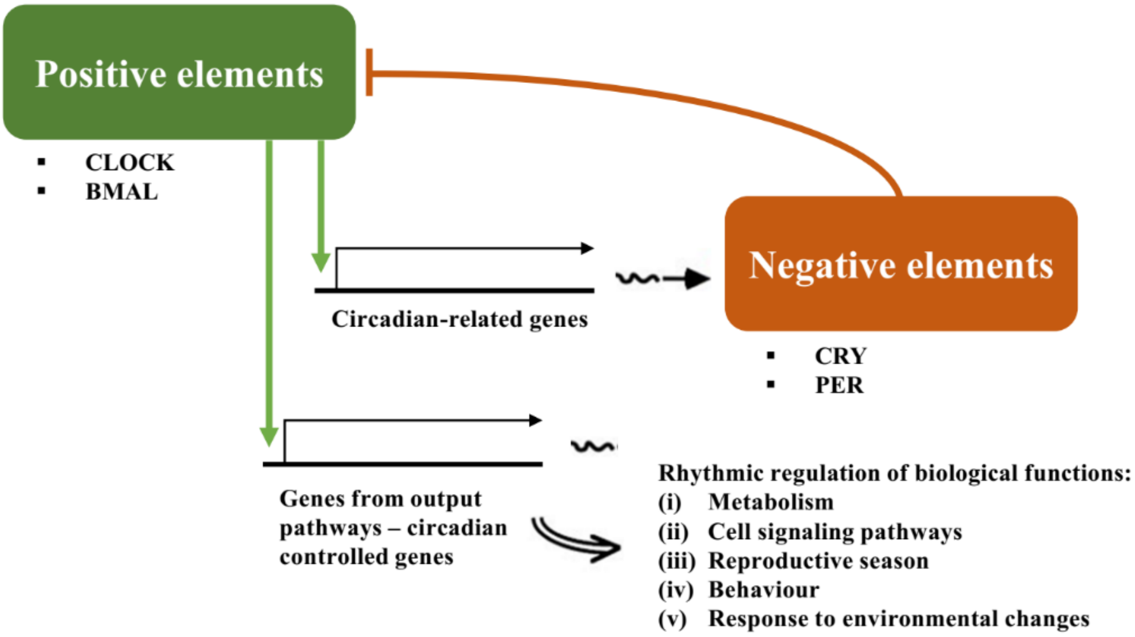


**Fig. S1.** Overview of the core circadian system and output pathways (adapted from Dunlap, 1999 [1])

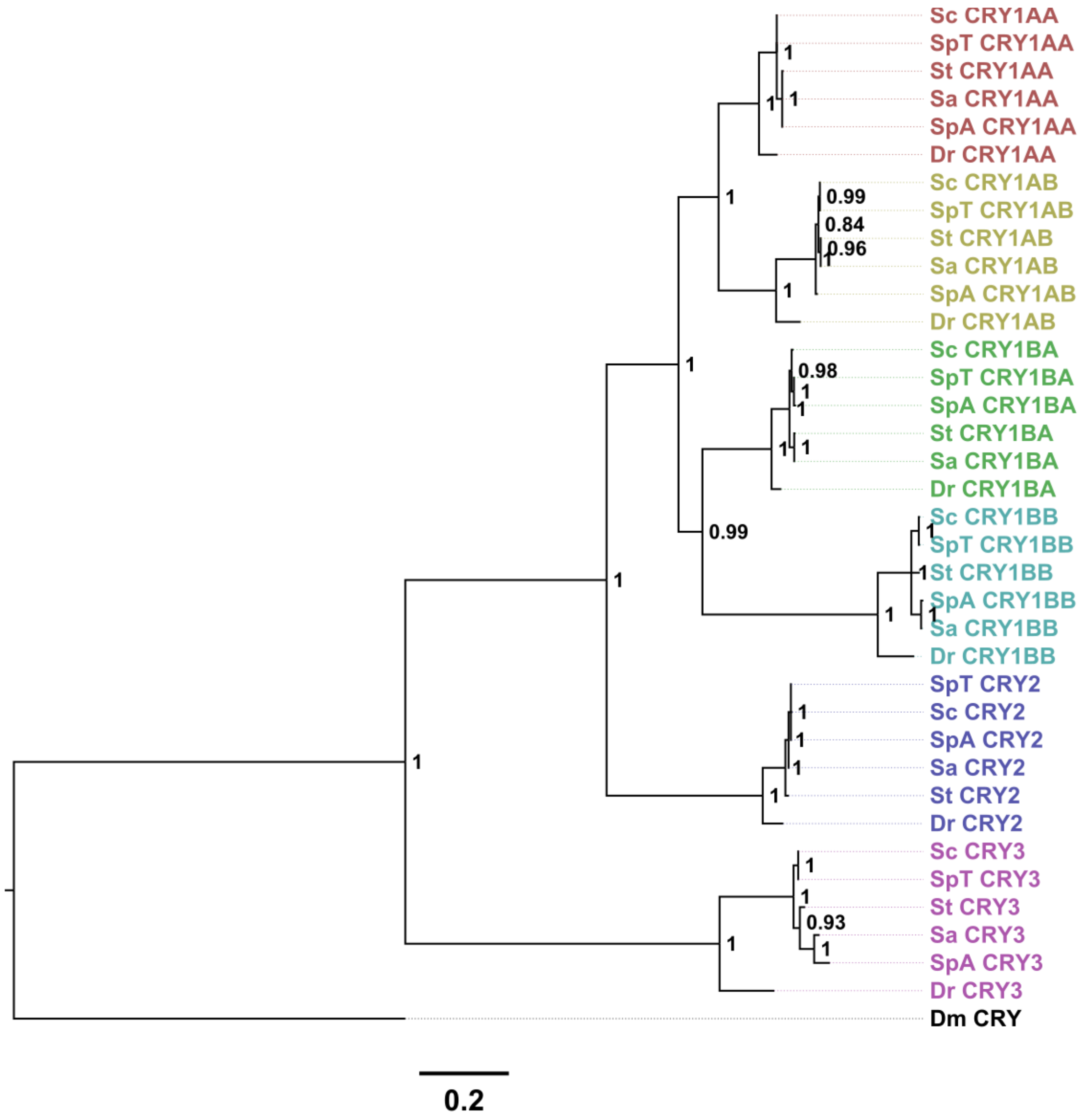
Sc CRY1AA
SpT CRY1AA
St CRY1AA
Sa CRY1AA
SpA CRY1AA
Dr CRY1AA
Sc CRY1AB
SpT CRY1AB
St CRY1AB
Sa CRY1AB
SpA CRY1AB
Dr CRY1AB
Sc CRY1BA
SpT CRY1BA
SpA CRY1BA
St CRY1BA
Sa CRY1BA
Dr CRY1BA
Sc CRY1BB
SpT CRY1BB
St CRY1BB
SpA CRY1BB
Sa CRY1BB
Dr CRY1BB
SpT CRY2
Sc CRY2
SpA CRY2
Sa CRY2
St CRY2
Dr CRY2
Sc CRY3
SpT CRY3
St CRY3
Sa CRY3
SpA CRY3
Dr CRY3
Dm CRY
0.99
0.84
0.96
0.98
0.99
0.93
0.2

**Fig. S2 (a)**

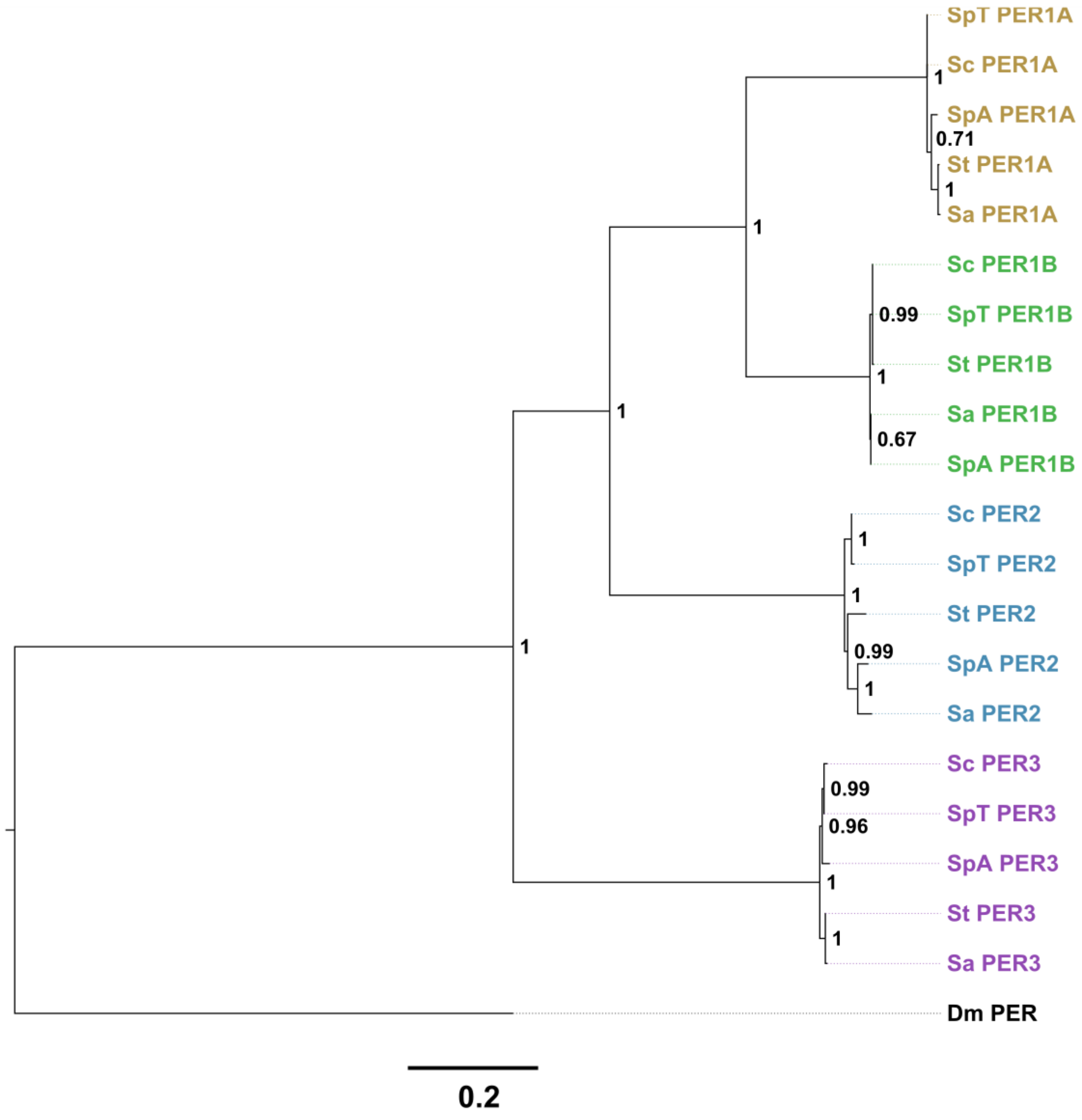

SpT PER1A
Sc PER1A
SpA PER1A
St PER1A
Sa PER1A
Sc PER1B
SpT PER1B
St PER1B
Sa PER1B
SpA PER1B
Sc PER2
SpT PER2
St PER2
SpA PER2
Sa PER2
Sc PER3
SpT PER3
SpA PER3
St PER3
Sa PER3
Dm PER
1
0.71
1
1
0.99
1
0.67
1
1
0.99
1
1
0.99
0.96
1
1
1
0.2


**Fig. S2 (b)**

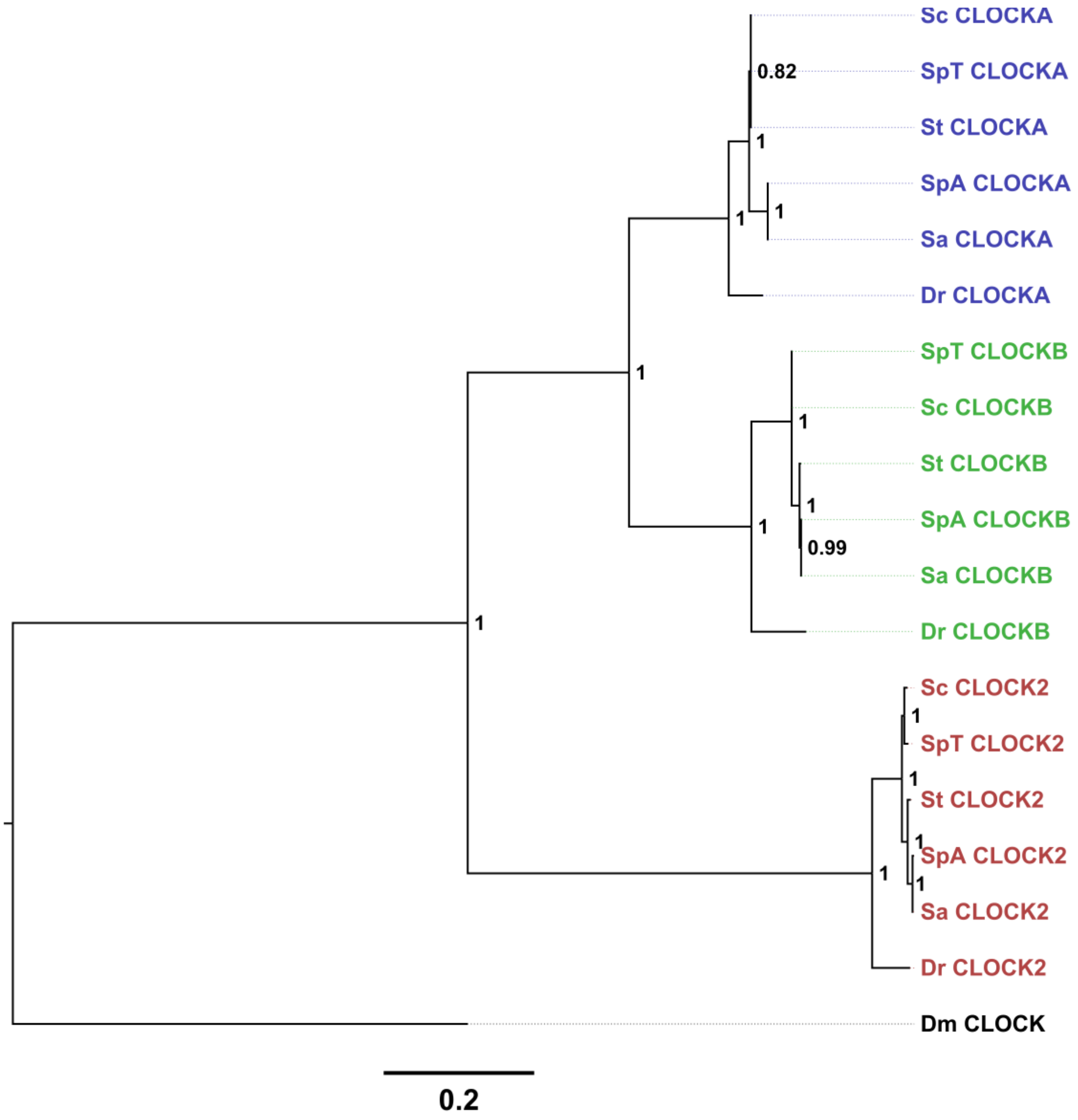
Sc CLOCKA
SpT CLOCKA
St CLOCKA
SpA CLOCKA
Sa CLOCKA
Dr CLOCKA
SpT CLOCKB
Sc CLOCKB
St CLOCKB
SpA CLOCKB
Sa CLOCKB
Dr CLOCKB
Sc CLOCK2
SpT CLOCK2
St CLOCK2
SpA CLOCK2
Sa CLOCK2
Dr CLOCK2
Dm CLOCK
0.82
0.99
0.2

**Fig. S2 (c)**

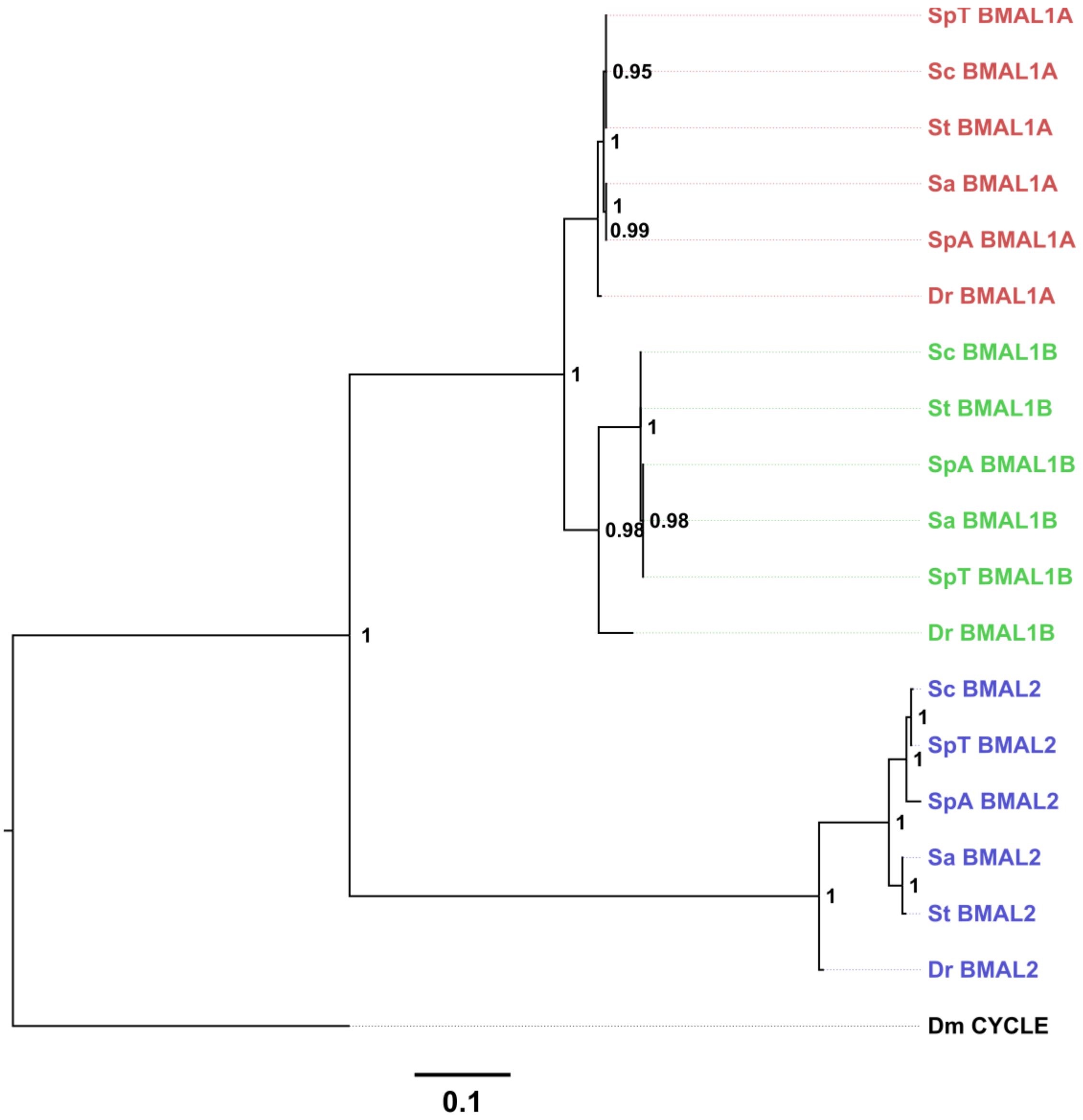
SpT BMAL1A
Sc BMAL1A
St BMAL1A
Sa BMAL1A
SpA BMAL1A
Dr BMAL1A
Sc BMAL1B
St BMAL1B
SpA BMAL1B
Sa BMAL1B
SpT BMAL1B
Dr BMAL1B
Sc BMAL2
SpT BMAL2
SpA BMAL2
Sa BMAL2
St BMAL2
Dr BMAL2
Dm CYCLE
0.95
1
1
0.99
1
1
0.98
0.98
1
1
1
1
1
1
0.1

**Fig. S2 (d)**

**Fig. S2.** A phylogenetic tree constructed by the Bayesian Inference method for **(a)** CRY proteins with fly (*Drosophila melanogaster*) CRY as outgroup using the LG substitution model [2] with a discrete Gamma distribution (+G) with 5 rate categories; **(b)** PER proteins with fly PER as outgroup using the JTT substitution model [3] with a discrete Gamma distribution (+G) with 5 rate categories and empirical amino acid frequencies from the data (+F); **(c)** CLOCK proteins with fly CLOCK protein as outgroup using JTT substitution model [3] using a discrete Gamma distribution (+G) with 5 rate categories and empirical amino acid frequencies from the data (+F); **(d)** BMAL proteins with fly CYCLE protein as outgroup using JTT substitution model [3] using a discrete Gamma distribution (+G) with 3 rate categories. Values on branch nodes represent Bayesian posterior probabilities. Sc, *Squalius carolitertii*; SpT, *Squalius pyrenaicus* (Tagus population); SpA, *Squalius pyrenaicus* (Almargem); St, *Squalius torgalensis*; Sa, *Squalius aradensis*; population); Dr, *Danio rerio*; Dm, *Drosophila melanogaster*.

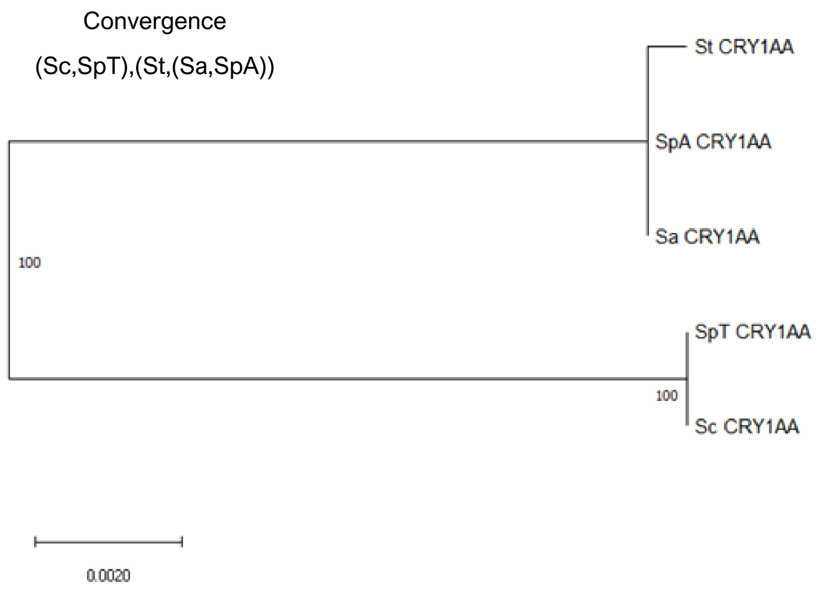


**Fig. S3 (a)**

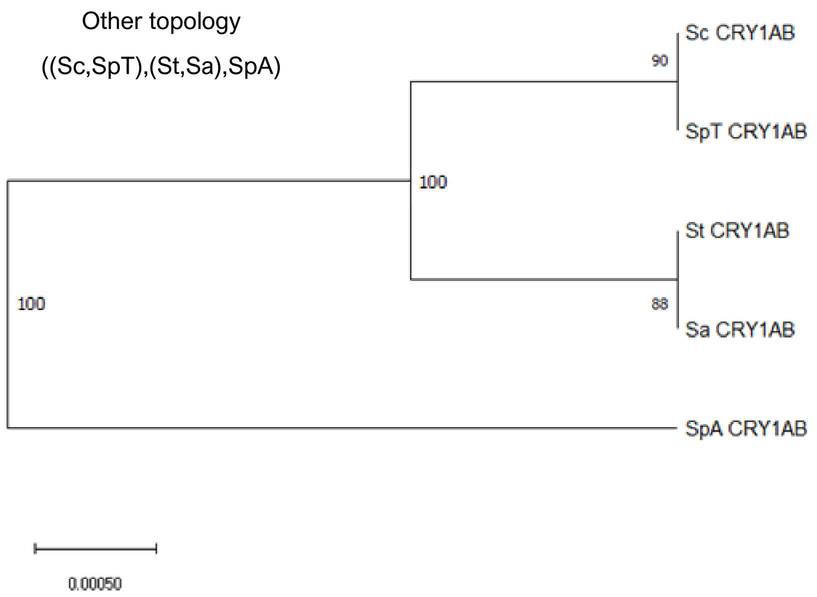


**Fig. S3 (b)**

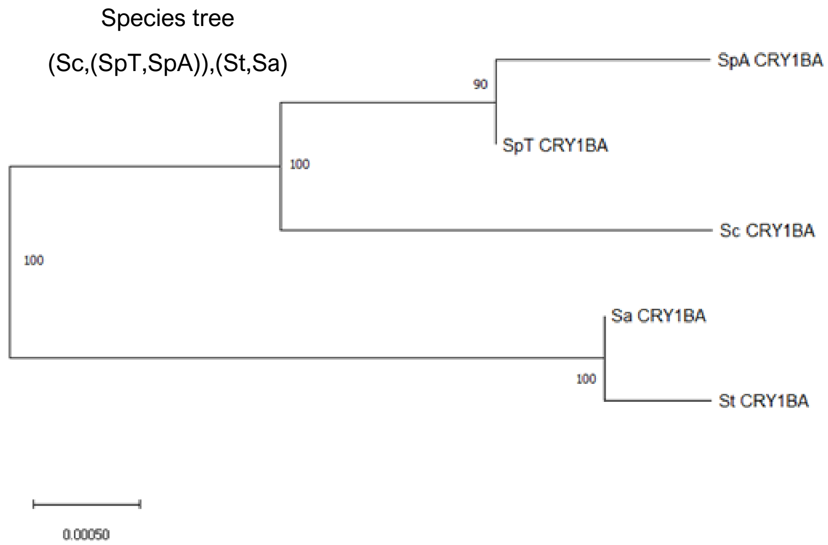


**Fig. S3 (c)**

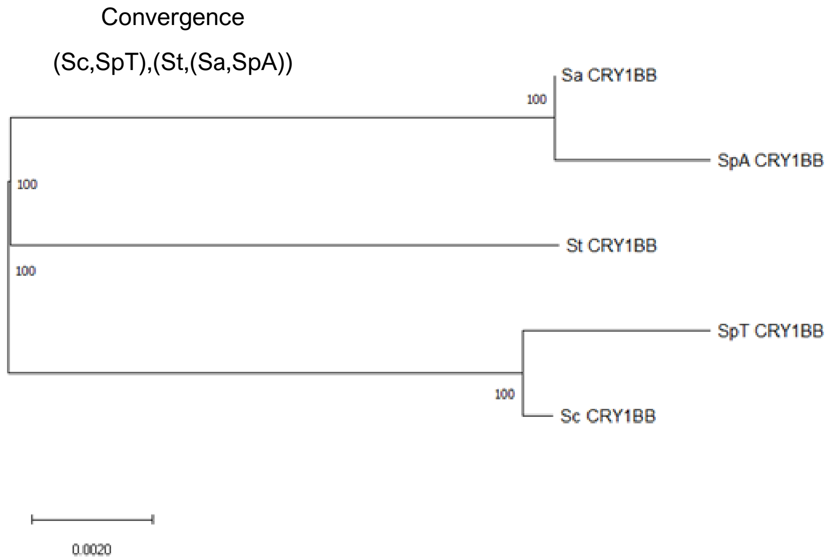


**Fig. S3 (d)**

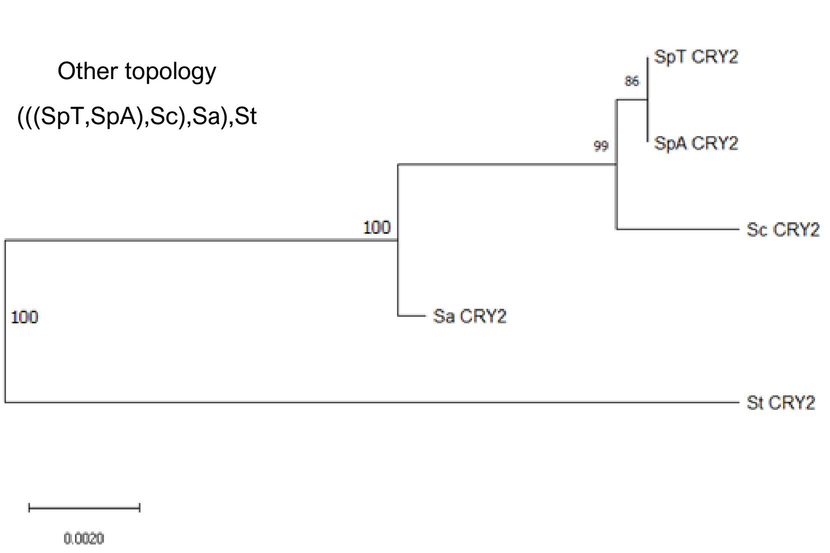


**Fig. S3 (e)**

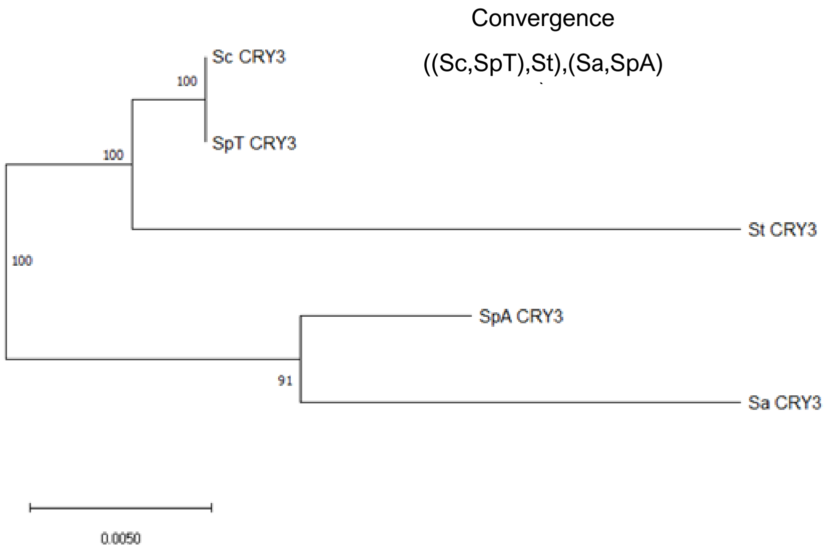


**Fig. S3 (f)**

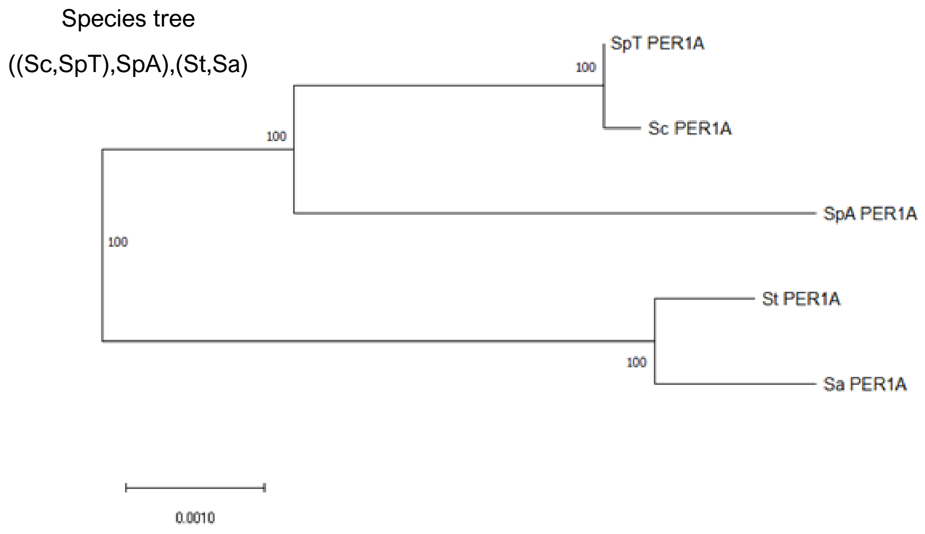


**Fig. S3 (g)**

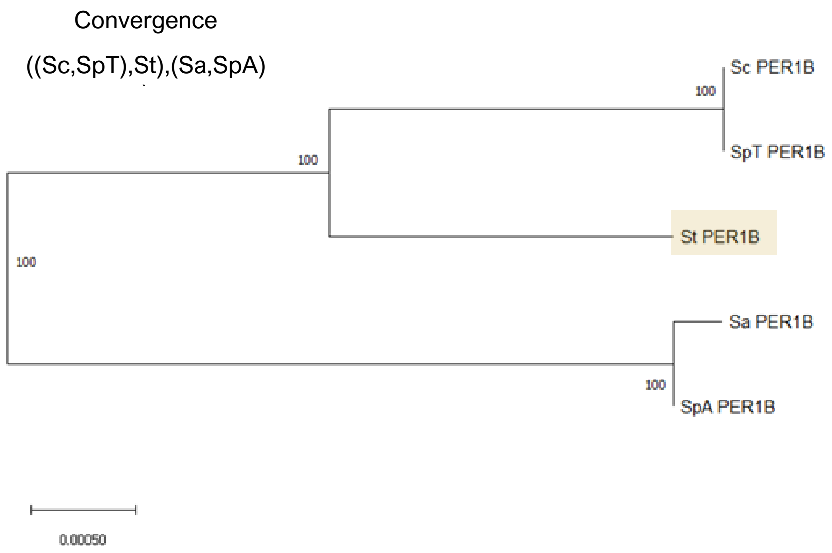


**Fig. S3 (h)**

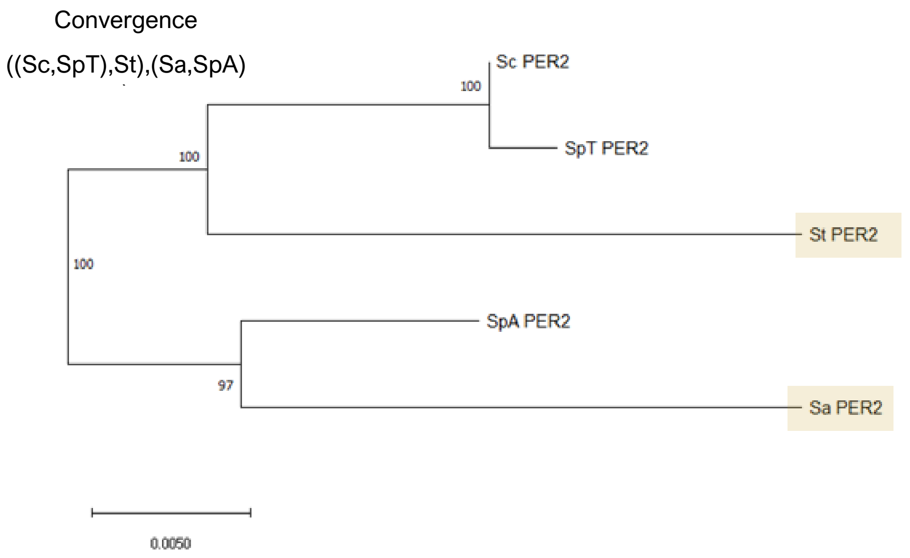


Fig. S3 (i)

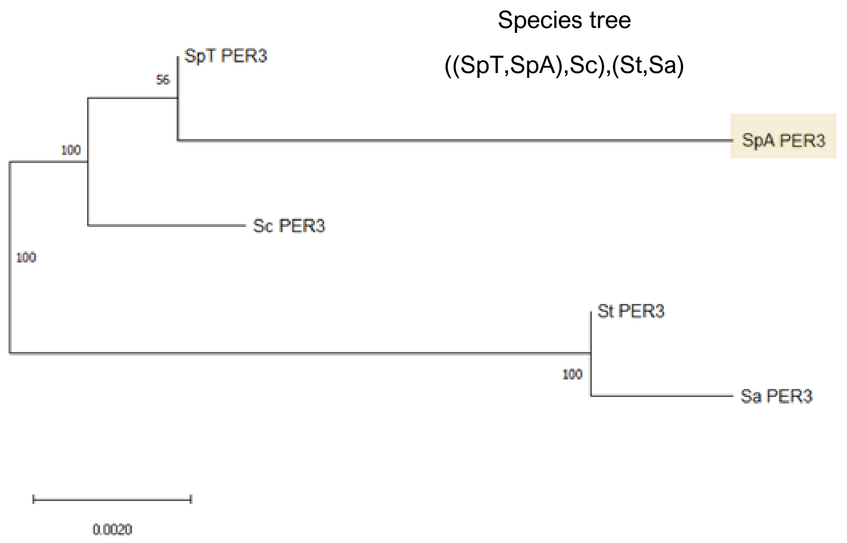


Fig. S3 (j)

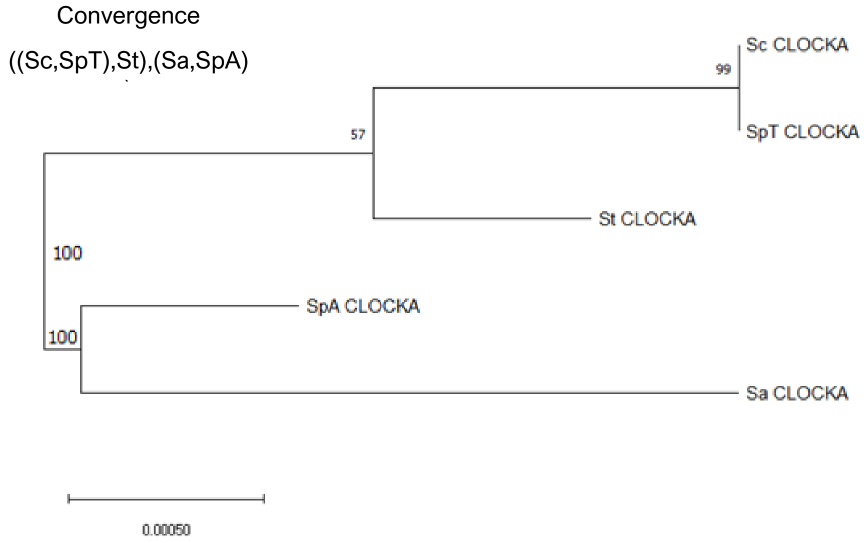


Fig. S3 (k)

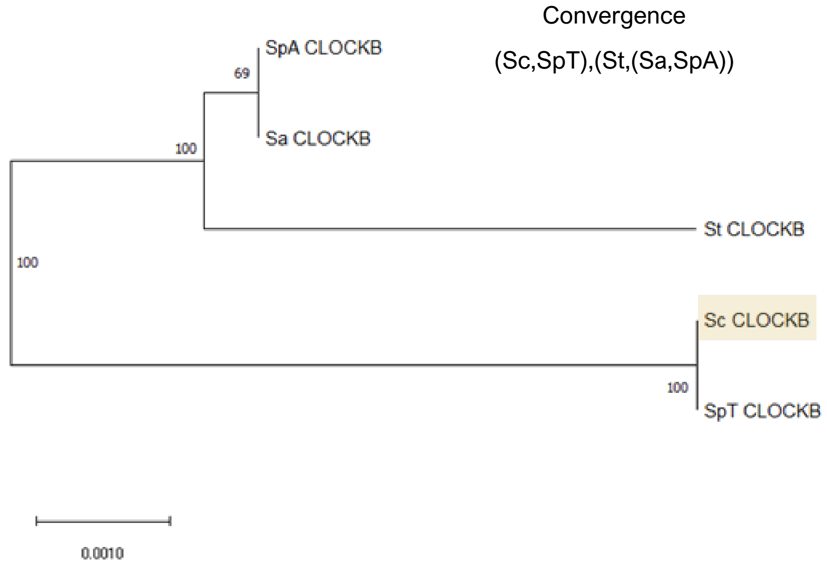


Fig. S3 (l)

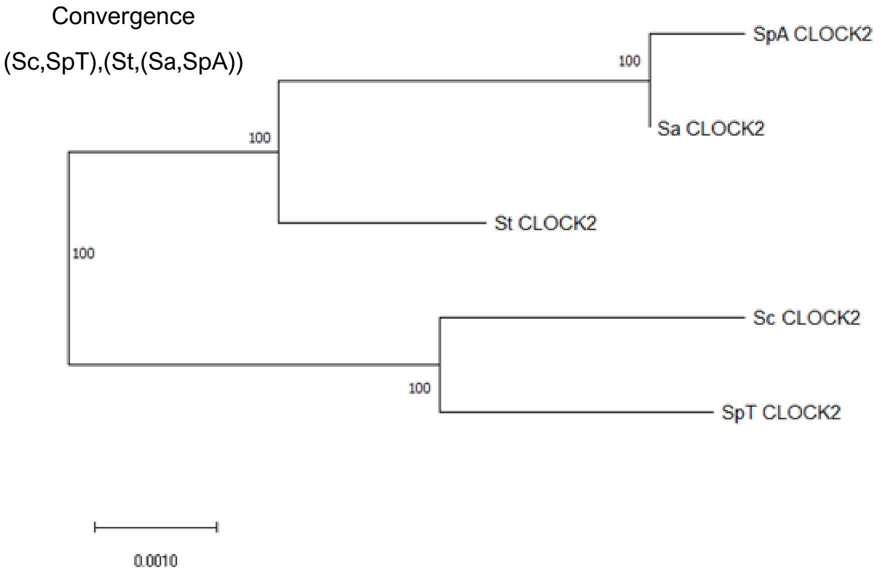


**Fig. S3 (m)**

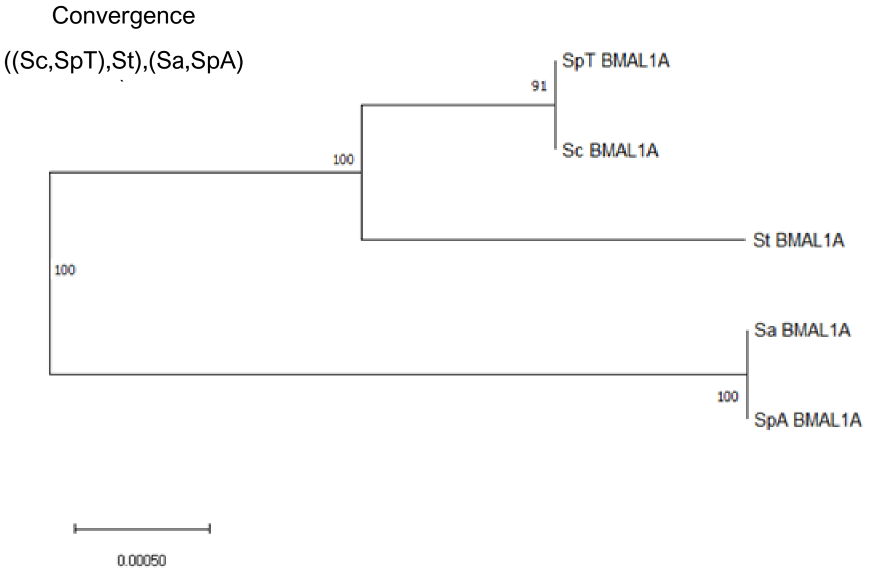


**Fig. S3 (n)**

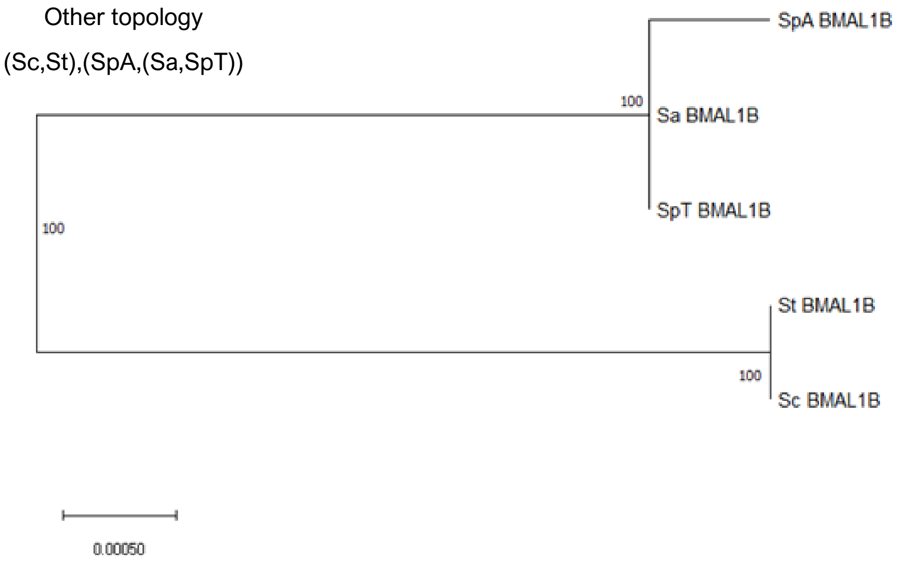


**Fig. S3 (o)**

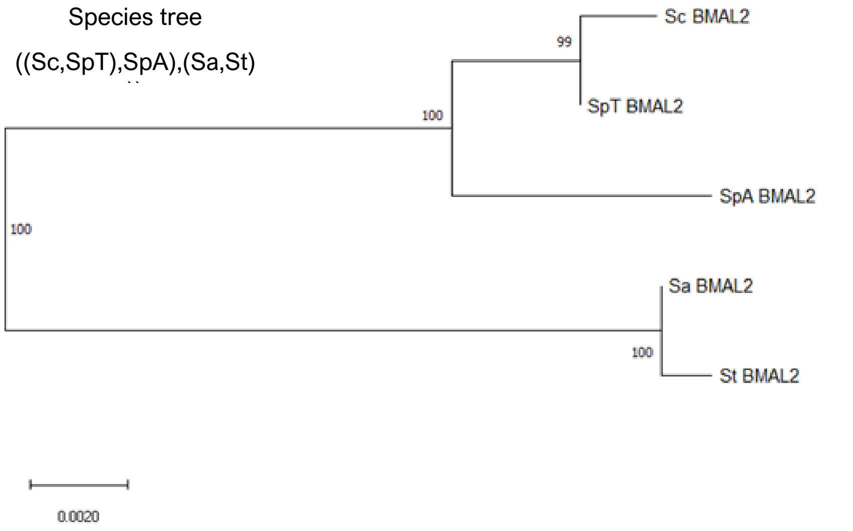


**Fig. S3 (p)**

**Fig. S3.** Unrooted maximum-likelihood phylogenetic trees for **(a)** *cry1aa,* **(b)** *cry1ab,* **(c)** *cry1ba,* **(d)** *cry1bb,* **(e)** *cry2,* **(f)** *cry3,* **(g)** *per1a,* **(h)** *per1b,* **(i)** *per2,* **(j)** *per3,* **(k)** *clocka,* **(l)** *clockb,* **(m)** *clock2,* **(n)** *bmal1a,* **(o)** *bmal1b,* **(p)** *bmal2.* Values on branch nodes represent bootstrap probabilities values based on 5000 replicates. Sc, *Squalius carolitertii;* SpT, *Squalius pyrenaicus* (Tagus population)*;* St, *Squalius torgalensis;* Sa, *Squalius aradensis;* SpA, *Squalius pyrenaicus* (Almargem population). Taxa highlighted in yellow correspond to species under positive selection on the branch-site analysis (see Table 2 and Table S5 for more information).

### SUPPLEMENTARY TABLES

**Table S1:** Circadian related genes identified with respective annotations obtained in functional annotation analysis. ENA accession numbers are for non-redundant *Squalius* sequences obtained by Sanger sequencing in this work.

*(Included in a separated excel file)*

**Table S2:** List of *Danio rerio* and *Drosophila melanogaster* Uniprot accession ID for target proteins and ENA accession IDs for corresponding coding genes.

*(Included in a separated excel file)*

**Table S3:** Patterns of protein-protein interactions for circadian-related protein predicted with STRING v10.5 with a threshold of 0.7 for score. Rows shaded in orange are highlighted proteins related to temperature responses; in blue are highlighted putative circadian proteins with secondary functions; in green are highlighted UV-induced DNA damage repairing proteins.

*(Included in a separated excel file)*

**Table S4:** Summary of gene-wide positive selection analysis in circadian-related genes using the BUSTED method implemented in Datamonkey webserver. A threshold of 0.1 was used for statistical significance (p-value<0.1). Rows shaded in grey correspond to genes whose test for positive selection was statistically significant.

*(Included in a separated excel file)*

**Table S5:** Summary of branch-site positive selection analysis using the aBSREL method implemented in Datamonkey webserver. Positive selection was only tested at the tips of the phylogeny and species are grouped in a single branch when there are not differences in their sequence. A threshold of 0.1 was used for statistical significance (p-value<0.1). Rows shaded in grey correspond to results whose test for positive selection was statistically significant.

*(Included in a separated excel file)*

**Table S6:** Summary of the results obtained by FEL analysis for pervasive negative selection in coding genes for circadian-related proteins. A threshold of 0.1 was assumed for significance level (p-value<0.1). Rows shaded in grey correspond to sites located in functional domains/motifs of the protein.

*(Included in a separated excel file)*

**Table S7:** Predicted physicochemical parameters (AI – Aliphatic index and pI – isolectric point) for each predicted protein. Each value represents the mean values for each parameter of each population (n=5). Different shades of white to grey refer to values of protein parameters that are statistically similar. The superscripts *a* to *j* indicate the comparisons that were statistically significant (p-value<0.05).

*(Included in a separated excel file)*

**Table S8:** List of primer pairs and respective sequences used in PCR to (re)sequence circadian-related genes with Sanger method in *Squalius* species.

*(Included in a separated excel file)*

**Table S9:** PCR conditions for each pair of primers (supplementary table S10) used in amplification of circadian-related genes.

*(Included in a separated excel file)*

### SUPPLEMENTARY REFERENCES

1. Dunlap JC. Molecular Bases for Circadian Clocks. Cell. 1999;96:271–290.

2. Le SQ, Gascuel O. An Improved General Amino Acid Replacement Matrix. Molecular Biology and Evolution. 2008;25:1307–20.

3. Jones DT, Taylor WR, Thornton JM. The rapid generation of mutation data matrices from protein sequences. Bioinformatics. 1992;8:275–82.
